# Separable roles for Microprocessor and its cofactors ERH and SAFB1/2 during microRNA cluster assistance

**DOI:** 10.1101/2025.09.09.675111

**Authors:** Renfu Shang, Niko Popitsch, Seungjae Lee, Stefan L. Ameres, Eric C. Lai

## Abstract

While most conserved microRNA (miRNA) transcripts harbor a suite of features that mediate their efficient biogenesis into small RNAs, some loci bear suboptimal attributes that enable additional layers of processing regulation. A notable example is cluster assistance, whereby a miRNA hairpin with suboptimal nuclear biogenesis can be enhanced by an optimal neighbor. This process involves local transfer of the Microprocessor complex, composed of the RNase III enzyme Drosha and its partner DGCR8, in concert with cofactors such as ERH and SAFB1/2. However, the mechanism(s) that underlie miRNA cluster assistance remain largely unclear. Here, we gain insights into this process by integrating mutant cells of Microprocessor and its cofactors with analysis of miRNA structure-function variants, biochemical tests and genomewide profiling. We define features of suboptimal miRNAs that render them dependent on cluster assistance, and distinguish amongst a network of proposed interactions amongst Microprocessor and its cofactors, to reveal a subset that are critical for cluster assistance. Most importantly, we use epistatic tests to separate and order the functional requirements for ERH and SAFB1/2 into a pathway. Our data indicate that ERH may engage in the process of Microprocessor transfer between hairpins, while SAFB factors (especially SAFB2) mediate recognition and stable binding of a suboptimal miRNA hairpin after Microprocessor transfer. Finally, we show how cluster assistance integrates into a feedback regulatory loop on Microprocessor, via Drosha-mediated cleavage of a suboptimal miRNA hairpin in the DGCR8 transcript. Altogether, our findings reveal complex regulatory transactions during biogenesis of clustered miRNAs.

## Introduction

Since the discovery of the first microRNA (miRNA) lin-4 and its critical regulatory target *lin-14* by Ambros and Ruvkun in 1993 (Lee et al. 1993; Wightman et al. 1993), a large community of researchers continues to study how miRNAs are generated, how they regulate gene expression, and how they impact biology. The bulk of well-expressed miRNAs in animals are generated by a canonical biogenesis pathway. In short, following the transcription of primary miRNAs (pri-miRNAs), the nuclear Microprocessor complex (composed of the RNase III enzyme Drosha and its partner DGCR8) recognizes and cleaves the hairpins to liberate the pre-miRNAs. Then pre-miRNA hairpins are exported to the cytoplasm and cleaved by the RNase III enzyme Dicer yielding ∼22 nucleotide (nt) miRNA duplexes. These duplexes are loaded into Argonaute proteins and a single strand is preferentially retained in the mature complex, which guides it to appropriate targets (Bartel 2018; Shang et al. 2023). As animal miRNA targets require only limited complementarity (∼7 nt) to positions 2-8 of the miRNA (Lai and Posakony 1997; Lai et al. 1998; Lai 2002; Brennecke et al. 2005), referred to as the seed region (Lewis et al. 2003; Bartel 2018), miRNAs tend to mediate extensive gene regulatory networks in metazoan species (McGeary et al. 2019).

As the gatekeeper for the canonical miRNA pathway, the Microprocessor complex must specifically select desired pri-miRNA substrates for efficient processing, while rejecting a sea of other plausible RNA hairpins. Biochemical and genomic studies reveal that optimal pri-miRNA hairpins harbor single-stranded flanking regions, a double-stranded stem of ∼35 basepairs (bps), a single-stranded terminal loop of >10 nts, and one or more sequence motifs within or close to the pri-miRNA hairpins (Zeng and Cullen 2005; Zeng et al. 2005; Han et al. 2006; Zhang and Zeng 2010; Auyeung et al. 2013; Ma et al. 2013; Fang and Bartel 2015; Nguyen et al. 2015; Kwon et al. 2016; Kwon et al. 2019). Recent cryo-EM structural studies of Microprocessor and its substrates provide insights into how such features are recognized (Jin et al. 2020; Partin et al. 2020; Garg et al. 2024), although the full picture of their interactions is still unclear.

Although one might imagine that conserved miRNAs have been honed by evolution to acquire optimal biogenesis features, rarely does any biochemical or biological process run at full steam, all of the time. For example, most canonical optimal miRNA hairpins harbor only a subset of the “menu” of positive biogenesis features (Fang and Bartel 2015; Kwon et al. 2019; Lee et al. 2023), which potentially enables regulation of processing. Perhaps more unexpected are miRNA loci that explicitly violate attributes of core canonical miRNA. In some cases, such highly suboptimal miRNA structures permit such substrates to escape the supervision of Microprocessor, and exploit other ribonucleolytic strategies to access the miRNA pathway. For example, mirtron hairpins lack optimal dsRNA stems and ssRNA flanking regions, but are instead cleaved by the splicing machinery(Okamura et al. 2007; Ruby et al. 2007). Transcription start site miRNA hairpins exemplified by the *mir-320* family lack optimal dsRNA stems and terminal loops, but their short hairpins are directly transcribed by RNA Polymerase II, prior to Dicer processing (Xie et al. 2013; Zamudio et al. 2014). Additional RNA processing strategies can yield functional miRNAs via Drosha- or Dicer-independent mechanisms (Yang and Lai 2011), and in fact it is possible to efficiently generate mammalian miRNAs independently of RNase III enzymes (Maurin et al. 2012).

Unexpectedly, the processing of some suboptimal miRNA hairpins still depends on Microprocessor, including Dicer-independent *mir-451* that bears a short stem and small terminal loop (Cheloufi et al. 2010; Cifuentes et al. 2010; Yang et al. 2010). How do such suboptimal hairpins access Microprocessor, when it is designed to exclude such substrates? Our group and others uncovered a mechanism termed cluster assistance, which underlies the nuclear processing of Microprocessor-dependent suboptimal miRNA hairpins (Fang and Bartel 2020; Hutter et al. 2020; Shang et al. 2020). The critical insight is that intrinsically suboptimal nuclear biogenesis of a miRNA hairpin can be enhanced by an optimal miRNA hairpin within the same primary transcript. Mechanistic tests support that this involves ordered recruitment of Microprocessor to the optimal hairpin and local transfer to the suboptimal hairpin (Shang et al. 2020).

While the “recruit-transfer” model is focused on Microprocessor, several trans-acting factors also promote miRNA cluster assistance (Fang and Bartel 2020; Hutter et al. 2020). An elegant genetic screen took advantage of the *mir-15a/16* cluster, where *mir-16* bears optimal structure but *mir-15a* is suboptimal with a short lower stem. The use of dual miRNA reporters for both members enabled CRISPR-Cas9 screening to distinguish between hits that affect both miRNAs (including core factors in miRNA biogenesis and target regulation) from ones that might specifically aid suboptimal miRNA biogenesis (and thus score only with the miR-15a reporter). This revealed how Scaffold Attachment Factor B1 and B2 (SAFB1/2) and Enhancer of Rudimentary Homolog (ERH) selectively promote biogenesis of suboptimal miRNA hairpins (Hutter et al. 2020). Biochemical analysis of Microprocessor associated proteins revealed that ERH binds directly to DGCR8 and participates in cluster assistance (Fang and Bartel 2020; Kwon et al. 2020).

Since this trio of initial studies, the DGCR8-ERH complex was solved (Kwon et al. 2020) and additional parameters of miRNA clusters assistance were enumerated (Shang and Lai 2023). Nevertheless, the mechanistic roles of these Microprocessor cofactors remain largely unclear. For example, it has not yet been distinguished if they facilitate Microprocessor transfer between hairpins, stabilize or enhance cleavage activity of Microprocessor on suboptimal miRNA hairpins, or facilitate higher order assemblies of Microprocessor. Moreover, in all of these scenarios, it remains unclear if ERH and SAFB1/2 all work together in a complex with DGCR8/Drosha, or if there might be separable roles of these cofactors. Thus, current reviews of miRNA cluster assistance envision that numerous possibilities are plausible (Shang et al. 2023; Kim et al. 2025) (**Supplementary Figure 1**).

Further insights are complicated by the fact that both ERH and SAFB1/2 have numerous, and still not fully defined, roles beyond miRNA biogenesis. For example, ERH is one of the most highly interconnected factors amongst RNA binding protein (RBP) complexes (Street et al. 2024), and SAFB1/2 are paralogous proteins with dual DNA-binding and RNA-binding architecture (Norman et al. 2016). Accordingly, simple profiling of mutant conditions will inevitably reflect a complex mixture of direct regulatory effects, of which miRNA cluster assistance is only one, along with a suite of indirect consequences. Instead, we pair genetic mutants with a series of biochemical and genomic profilings, along with panels of structure-function variants, allowing us to specifically probe the requirements of Microprocessor cofactors and different steps of nuclear miRNA biogenesis. We find complex roles for these factors to support Microprocessor activity, including distinct effects on different types of suboptimal miRNAs, and separable roles for ERH and SAFB2 during Microprocessor recruitment and transfer.

## Results

### Both ERH and SAFB2 are required for efficient processing of suboptimal *mir-451*

To study the roles of ERH and SAFB1/2 in miRNA cluster assistance, we generated a collection of mutant HEK293T cells. While we succeeded in recovering protein-null *SAFB2* mutants, following multiple rounds of engineering, we found that all *ERH* and *SAFB1* mutant clones retained a wildtype allele (**Supplementary Figure 2**). We conclude that SAFB1 and ERH are both cell-essential in HEK293T cells. Since multiple core miRNA factors can be fully deleted from HEK293T cells (Bogerd et al. 2014; Aguado et al. 2017), this indicates that ERH and SAFB1 must harbor non-miRNA functions. Accordingly, we further depleted these factors using *ERH-RNAi* in *ERH[+/-]* cells and *SAFB1-RNAi* in *SAFB1[+/-], SAFB2[-/-]* cells (**Figure 1A**). We also made a triple mutant by knocking down ERH and SAFB1 in *ERH[+/-]; SAFB1[+/-], SAFB2[- /-]* cells. These represent the strongest individual and combined depletions that we could obtain for these cofactors. Nevertheless, a “silver lining” to these efforts was that cells were not adapted to complete knockout of these factors, which is an advantage to the study of these models.

**Figure 1.**
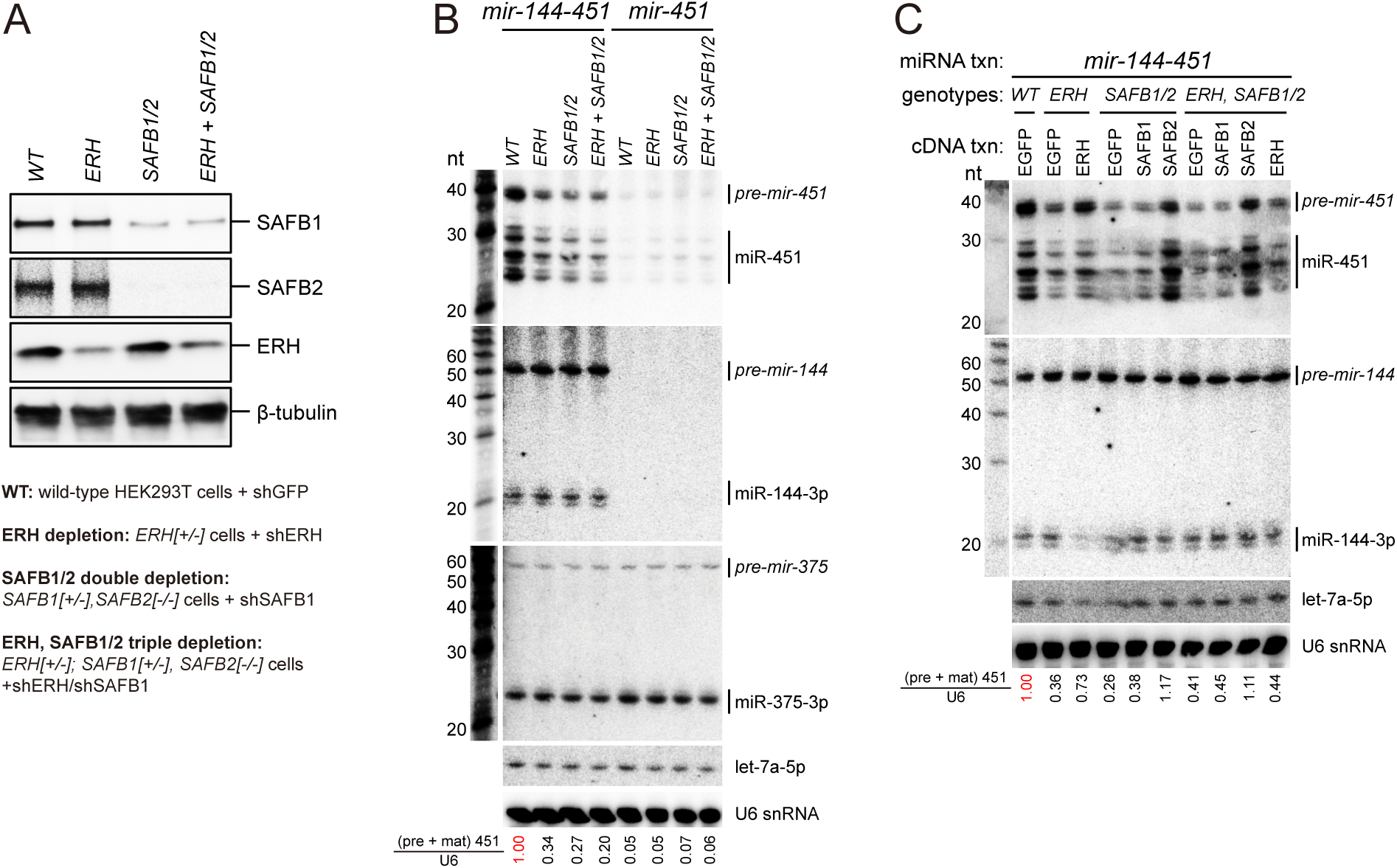
Biogenesis of suboptimal *mir-451* requires cluster assistance from ERH and SAFB1/2. (A) Deletion and/or depletion of *ERH* and *SAFB1/2* in HEK293T cells. *ERH* mutant cells were generated by shRNA-mediated knockdown in an *ERH[+/-]* heterozygous cell line. SAFB1/2 double mutant cells were generated by knockdown of *SAFB1* in a *SAFB1[+/-], SAFB2[-/-]* cell line. *ERH+SAFB1/2* triple mutant cells were generated by knockdown of *ERH* and *SAFB1* in an *ERH[+/-]; SAFB1[+/-], SAFB2[-/-]* cell line. Western blot validates strong loss of ERH and SAFB1 proteins and absence of SAFB2 in the respective mutant genotypes. β-tubulin was probed as control. (B) Northern blotting to assay processing of *mir-144/451* cluster or solo *mir-451* in wildtype or cofactor depleted HEK293T cells. Co-transfected *mir-375* and endogenous let-7a and U6 snRNAs were probed as controls. RNA size markers (nt) are shown at left. Microprocessing of suboptimal *pri-mir-451*, but not optimal *pri-mir-144* and *pri-mir-375*, is significantly impaired in *ERH* and/or *SAFB1/2* mutant cells. However, lack of the neighboring optimal *mir-144* hairpin ablated *pri-mir-451* processing, even with all the cofactors present. Normalized quantifications of (pre + mature) miR-451 total levels to U6 snRNA levels using Multi Gauge V3.0 were shown below the gel. (C) Rescue of *pri-mir-451* processing in mutant cells. While ERH overexpression can restore *mir-451* biogenesis in ERH mutant cells, only SAFB2 overexpression restored *mir-451* biogenesis in *SAFB1/2* and *ERH+SAFB1/2* mutant cells.

We examined the biogenesis of a *mir-144/451* construct transfected into these cells. To note, miR-451 is a non-canonical miRNA that transits an unusual biogenesis pathway, yielding atypical banding pattern on Northern blots {Cheloufi, 2010 #6432}{Cifuentes, 2010 #6434}{Yang, 2010 #6820;Yang, 2012 #10331}. To summarize, cleavage of pri-mir-451 by Drosha/DGCR8 yields a 42 nt *pre-mir-451* hairpin that is too short to be cleaved by Dicer, and instead loads directly into Ago proteins. Of the four human Ago family members, only Ago2 is capable of cleaving its 3’ arm, yielding a 30 nt product. This is further 3’ trimmed by PARN {Yoda, 2013 #13489} into heterogeneous, active, miR-451 species, which differ from typical miRNA products that are typically a distinct species in the 21-24 nt range (**Figure 1B**).

Extending previous work on the dependency of miR-451 on ERH (Fang and Bartel 2020; Kwon et al. 2020), we found that maturation of suboptimal *mir-451* was substantially and selectively inhibited in *ERH* single, *SAFB1/2* double and *ERH/SAFB1/2* triple mutant cells (**Figure 1B**). In contrast, *mir-144* was unaffected by any of these mutations, and neither were co-transfected optimal *mir-375* or endogenous *let-7a* (**Figure 1B**). Notably, the nuclear processing of a solo *mir-451* construct, lacking its optimal *mir-144* hairpin neighbor, was strongly abrogated in wildtype cells, and was not further repressed by loss of ERH and/or SAFB1/2 (**Figure 1B**). We conclude that presence of the optimal miRNA hairpin is the first and most essential step for cluster assistance, and that cofactors opt into the following steps to enhance processing of the adjacent suboptimal hairpin.

SAFB1 and SAFB2 have similar domain organization and sequences. To evaluate the extent of their functional redundancy, we compared *SAFB1*-RNAi in a *SAFB1[+/-]* cell line with *SAFB2* null cells. Loss of *SAFB1* did not affect *mir-451* processing, while mutation of *SAFB2* inhibited miR-451 biogenesis to a comparable extent as *ERH* depletion (**Supplementary Figure 3A**). However, combined *SAFB1/2* loss had a stronger effect than *SAFB2* loss alone (**Supplementary Figure 3A**). These data mirror results with the *mir-15a/16* suboptimal miRNA cluster in murine Baf3 cells (Hutter et al. 2020). Thus, SAFB1/2 have overlapping roles in cluster assistance, but SAFB2 plays a dominant role in cell lines from different species. We emphasize that SAFB1 appears to be cell-essential in HEK293T, while protein-null SAFB2 lines were easily recovered (**Figure 1A**). Accordingly, SAFB1/2 paralogs must have some distinct functions, i.e. the preferred role of SAFB2 in cluster assistance is not solely due to, for example, its preferred expression in HEK293T cells.

Next, we conducted rescue assays in our panel of mutant cells, by transfecting cognate and non-cognate cluster assistance factors. Ectopic expression of all these factors rescued miR-451 biogenesis in their corresponding mutants, confirming that their defects were not due to unanticipated genetic aberrations. More interesting were the results from cross-rescue tests. Ectopic ERH could not rescue *SAFB1/2* mutants, suggesting that SAFB proteins exert a non-redundant role from ERH (**Supplementary Figure 3B**). As well, only SAFB2, but not SAFB1, rescued *mir-451* maturation in *SAFB1/2* double depleted cells (**Figure 1C**). This is consistent with our data that SAFB2 is more important than SAFB1 for cluster assistance in HEK293T cells. Unexpectedly, SAFB2 partially rescued miR-451 processing in both *ERH* mutants (**Supplementary Figure 3B**) as well as *ERH/SAFB1/2* triple mutant cells (**Figure 1C**). Such epistasis data hint that SAFB and ERH not only have distinct roles, but that SAFB2 may function downstream of ERH. These findings inform our subsequent mechanistic tests.

### Specific mutations of *mir-451* decouple the biogenesis requirements for ERH and SAFB2

Meta-analysis of small RNA profiling showed that a substantially overlapping set of miRNAs depend on both ERH and SAFB1/2 (Kwon et al. 2020), suggesting that they function in a common process underlying cluster assistance. Such a scenario might be ruled out if any miRNA was explicitly shown to depend on only one of these factors, but none have yet been characterized. Indeed, our tests of suboptimal *mir-451*, previously shown to depend on ERH, reveal that it also depends on SAFB1/2 (**Figure 1**).

Accordingly, we attempted to decouple their roles by introducing mutations into *mir-451*, building on our extensive engineering of the *mir-144* and *mir-451* hairpins (Shang et al. 2020; Shang et al. 2022; Shang and Lai 2023). We were guided by previous observations that suboptimal *mir-15a* hairpin has a short lower stem, while *mir-451* has a small terminal loop and a short upper stem. Thus, we designed *mir-451* mutants (within the context of the *mir-144/451* cluster) that lack one or more suboptimal hairpin features, namely a single-stranded apical loop of >10 nts, a double-stranded upper stem of ∼22 bp, and a stable lower stem of ∼11 bp (**Figure 2A-B**). As a control, we optimized the *pri-mir-451* hairpin (451LSM) to include all the above optimal structural features, which bypasses the need for an optimal miRNA neighbor (Shang et al. 2020).

**Figure 2.**
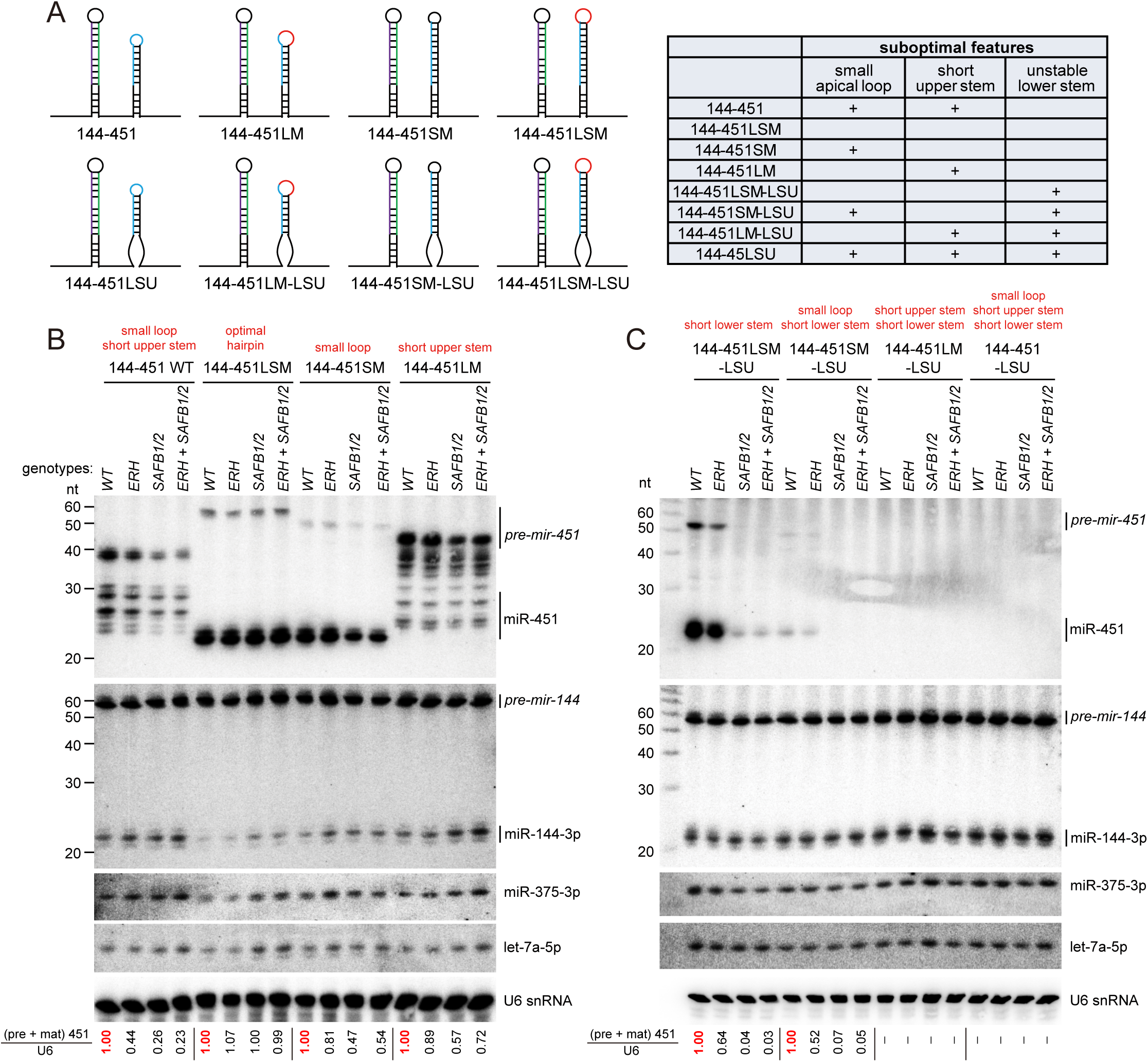
Engineered suboptimal miRNAs separate dependency on ERH and SAFB2. (A) Schematic of *mir-144/451* variants. We constructed an optimized *pri-mir-451* (451LSM) or variants with different suboptimal structural features, such as small apical loop, short upper stem, unstable lower stem or combinations of these features. (B-C) Northern blotting for the processing of *mir-144/451* variants with stable lower stem (B) or unstable lower stem (C) in WT, ERH and/or SAFB2 depleted cells. (B) Compared with wildtype *mir-451* containing both small apical loop and short upper stem, *mir-451* variants with only one of the two suboptimal features (451SM or 451LM) were less dependent on ERH and SAFB2. (C) A *mir-451* variant bearing only an unstable lower stem (451LSM-LSU) was especially dependent on SAFB factors, but only mildly dependent on ERH. When *mir-451* harbors an unstable lower stem and another suboptimal feature (451SM-LSU, 451LM-LSU or 451-LSU variants), its processing was fully suppressed, even when paired with an optimal *mir-144* neighbor and in the presence of all cluster assistance factors. The quantified ratios were normalized to the red lane in each *mir-451* mutant set.

We transfected *mir-451* mutants into wildtype and cofactor-depleted cells, and compared their biogenesis using Northern blotting. Consistent with our previous data, the optimized *mir-451* hairpin (451LSM) was independent of ERH and SAFB1/2 (**Figure 2B**). While the processing of *mir-451* mutants with either a small apical loop (451SM) or a short upper-stem (451LM) showed some dependence on ERH and SAFB2, their dependence was weaker than wildtype *pri-mir-451*, which harbors both small apical loop and short upper stem (**Figure 2B**). Since the presence of the neighboring optimal *mir-144* hairpin alone can significantly enhance the processing of *pri-mir-451SM* and *pri-mir-451LM* (Shang et al. 2020), the roles of ERH and SAFB2 in their processing are limited.

We next examined a *mir-451* mutant bearing an optimized loop, but harboring an unstable lower stem (451LSM-LSU). Strikingly, this mutant was only subtly dependent on ERH, but strongly dependent on SAFB1/2 (**Figure 2C**). This implies that SAFB2 is particularly critical when the miRNA hairpin lacks a stable lower stem. Importantly, the behavior of 451LSM-LSU showed that it is possible to decouple the requirements for the two types of cluster assistance factors. Finally, we mutated the *mir-451* lower stem in combination with another suboptimal feature, either small apical loop (451SM-LSU), short upper stem (451LM-LSU), or both (451-LSU) (**Figure 2A**). All of these variants fully blocked the biogenesis of miR-451, even in the presence of an optimal miRNA neighbor and with all endogenous cofactors intact (**Figure 2C**). Altogether, our tests on various *mir-451* mutants reveal that miRNAs with different suboptimal features exhibit selective dependence on ERH and SAFB2.

### ERH promotes cluster assistance, but this process does not require DGCR8/ERH interaction

With these data and reagents in hand, we proceeded to dissect the roles of Microprocessor cofactors, beginning with ERH. Human ERH is only 104 aa long, lacks an RNA binding domain, and does not interact with RNA directly. However, it is a constituent of numerous RBP complexes (Street et al. 2024) and associates with Microprocessor via DGCR8 (Kavanaugh et al. 2015; Fang and Bartel 2020; Kwon et al. 2020) (**Figure 3A**). Thus, we tested for interactions between ERH and miRNA pathway factors using co-immunoprecipitation (co-IP) assays. ERH interacted robustly with DGCR8 and weakly with Drosha (**Figure 3B**), consistent with prior studies. Co-IP assays with luciferase and Dicer served as negative controls.

**Figure 3.**
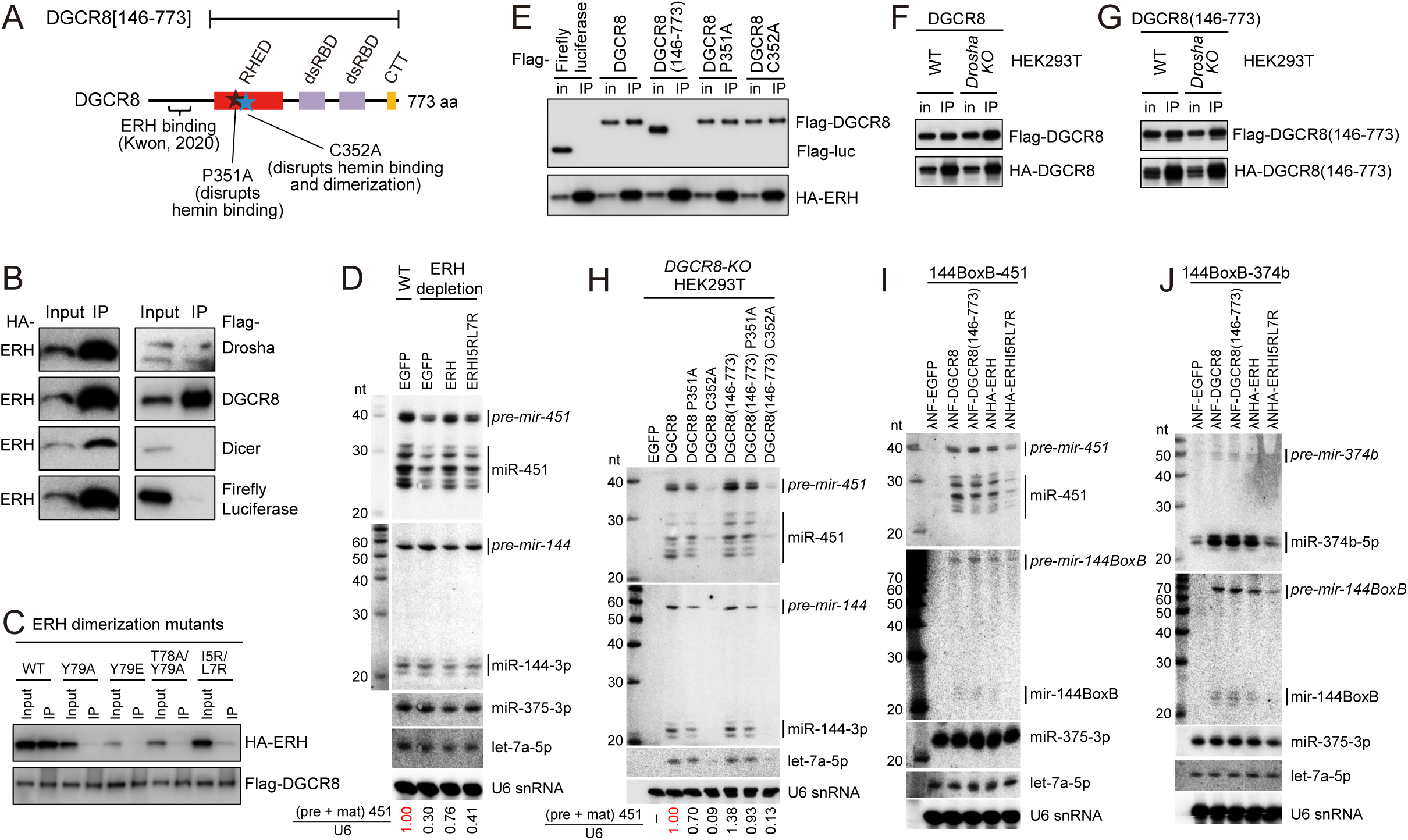
Interaction of ERH and DGCR8 is not required for cluster assistance. (A) Schematics of DGCR8 domains and protein/cofactor binding regions. (B) ERH interactions with miRNA pathway factors. Co-IP tests showed that ERH has strong interaction with DGCR8, moderate interaction with Drosha (likely indirect), and no interaction with Dicer or control Firefly luciferase. (C) Dimerization ability of ERH is required for its interaction with DGCR8. All dimerization mutants (Y79A, Y79E, T78A/Y79A and previously reported I5R/L7R) of ERH abrogated their interactions with DGCR8. (D) Northern blotting showed that ERH dimerization mutant (I5R/L7R) exhibited less efficient restoration on suboptimal *pri-mir-451* processing than wildtype ERH, indicating its dimerization is required for cluster assistance. (E) ERH interactions with wildtype and mutated DGCR8 by co-IP assays. N-terminally truncated DGCR8[146-773] fails to interact with ERH, while DGCR8 dimerization mutants (P351A and C352A) maintain interactions with ERH. (F-G) Both wildtype DGCR8 (F) and DGCR8[146-773] (G) can homodimerize, in both WT and *Drosha-KO* cells. Thus, ERH is not essential for the dimerization ability of DGCR8. (H) In a *DGCR8-KO* cell line (from Shuo Gu lab), both full-length and DGCR8[146-773] rescued *mir-144/451* processing. In fact, DGCR8[146-773] that cannot bind ERH provided stronger rescue for *pri-mir-451* processing than WT DGCR8. Meanwhile, DGCR8 dimerization mutants cannot (C352A) or only partially (P351A) restore *mir-144/451* processing. Thus, the interaction between ERH and DGCR8 is dispensable for cluster assistance. (I-J) Tethering assays for suboptimal *mir-451* (I) or *mir-374b* (J) confirm that the ERH-interacting region in DGCR8 is not essential for cluster assistance, but that DGCR8 dimerization is required.

Structural studies revealed how human ERH homodimerizes and interacts with the N-termini of DGCR8 dimers, forming a stable 2:2 complex (Kwon et al. 2020). To assess whether ERH homodimerization is required for interaction with DGCR8, we tested a previously reported ERH mutant (I5R/L7R) and generated additional mutants (Y79A, Y79E, T78A/Y79A) based on its structure (Kwon et al. 2020). We used pulldown assays to confirm that all of these lack homodimerization capacity (**Supplementary Figure 4A**). These ERH dimerization mutants almost completely fail to associate with DGCR8, indicating that the interaction interface of these factors requires ERH dimerization (**Figure 3C**). In a rescue assay, the ERH dimerization mutant (I5RL7R) did not enhance processing of suboptimal *mir-451* hairpin (**Figure 3D**).

These results clearly confirmed interactions between ERH and DGCR8, and that ERH requires dimerization to mediate miRNA cluster assistance. However, can we conclude from this that ERH:DGCR8 interaction is essential for cluster assistance? To test this, we truncated the N-terminus with DGCR8[146-773], which fully abrogates interaction with ERH (**Figure 3E**), consistent with published data (Kwon et al. 2020). In addition, we tested DGCR8 mutants that compromise hemin binding (P351A) (Barr et al. 2011; Barr et al. 2012) or lack both hemin-binding and dimerization abilities (C352A) (Faller et al. 2007). Importantly, both mutants can still interact with ERH (**Figure 3E**). Notably, truncated DGCR8[146-773] that cannot bind ERH, still homodimerized efficiently as wt DGCR8, in both wildtype and *Drosha* null cells (**Figure 3F-G**). Therefore, DGCR8 dimerization does not require DGCR8-ERH interaction or ERH dimerization.

Using these reagents, we assessed if the ERH-interacting domain in DGCR8 is required for cluster assistance. We expressed wildtype and variant DGCR8 proteins in *DGCR8* null cells (Dai et al. 2016), which accumulated to comparable levels (**Supplementary Figure 4B**), and compared their activities on *mir-144/451* processing. Surprisingly, DGCR8[146-773] completely rescued biogenesis of both miR-144 and miR-451 in *DGCR8-KO* cells, and even exhibited higher activity than wildtype DGCR8 to support biogenesis of suboptimal *mir-451* (**Figure 3H**). On the other hand, loss of DGCR8 dimerization ability, especially the C352A mutant, abrogated the processing of optimal *mir-144*, suboptimal *mir-451*, and endogenous *let-7a* hairpins (**Figure 3H**). This was the case regardless of whether the ERH-interacting domain was present or not. We repeated these tests in an independent *DGCR8-KO* cell line and obtained similar results (**Supplementary Figure 4C**).

We further tested the dependence on ERH-DGCR8 interaction for cluster assistance using our established λN-BoxB tethering assay (Shang et al. 2020). Direct recruitment of Microprocessor, containing full length or N-terminal truncated DGCR8, to the neighboring miRNA hairpin enhanced suboptimal *mir-451* processing (**Figure 3I**). Notably, tethering ERH to the optimal miRNA hairpin also enhanced *mir-451* biogenesis, but the dimerization and DGCR8 interacting mutants (ERHI5RL7R) could not do so (**Figure 3I**). We obtained similar results upon tethering the panel of ERH variants near another suboptimal miRNA, *mir-374b* (**Figure 3J**).

These tests support two key conclusions regarding cluster assistance. First, recruitment of Microprocessor to a miRNA cluster by tethering ERH, via interactions of ERH:DGCR8, can effectively promote suboptimal miRNA processing. However, if a sufficient level of Microprocessor is locally present, then the interaction between ERH and DGCR8 is dispensable for functional cluster assistance.

### Minimal SAFB2 promotes cluster assistance, but this process does not require Drosha/SAFB2 interaction

We next explored the mechanism of SAFB1/2 in miRNA cluster assistance. As SAFB1/2 are broadly involved in numerous cellular processes via multiple protein domains, we took advantage of SAFB2[512-726] (**Figure 4A**), a minimal separation of function fragment that is sufficient to promote miRNA cluster assistance (Hutter et al. 2020). We first validated that truncated SAFB2 could rescue suboptimal *mir-451* processing to a comparable extent as full-length SAFB2, in both *SAFB2-KO* and *SAFB1/2* double mutant cells (**Figure 4B-C**). Optimal miRNAs tested were not affected by truncated SAFB2, confirming that this minimal fragment has specific and sufficient roles in suboptimal miRNA processing (**Figure 4B-C**).

**Figure 4.**
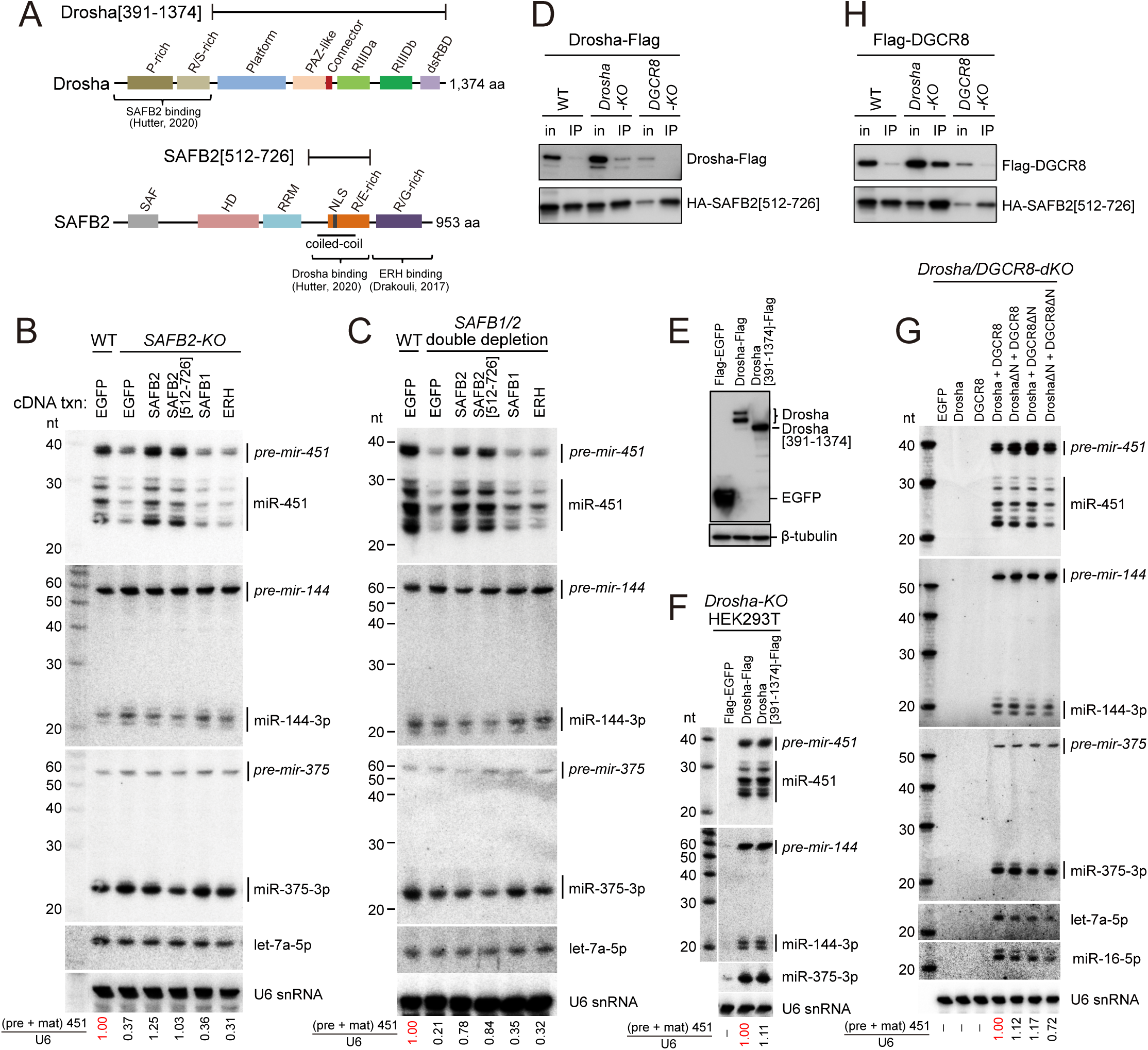
The N-terminal region of Drosha is dispensable for cluster assistance. (A) Schematics of Drosha and SAFB2 domains and protein binding regions. (B-C) A minimal fragment of SAFB2 containing only the RE-rich domain and a short upstream region can replace full length SAFB2 during suboptimal miRNA processing. In rescue assays of *SAFB2* single mutant (B) or *SAFB1/2* double mutant (C) HEK293T cells, SAFB2[512-726] restored nuclear cleavage of suboptimal *pri-mir-451* as efficiently as full length SAFB2. At the same time, ectopic SAFB1 and ERH were incapable of rescuing *mir-451* biogenesis in *SAFB2-KO* cells. (D) Co-IP tests showed very weak interactions between Drosha and functional active SAFB2[512-726], either in WT or Microprocessor null cells. (E) Expression of full length and N-terminal truncated Drosha in *Drosha-KO* HEK293T cells. (F) Restoration of *mir-144/451* and control *mir-375* processing in *Drosha-KO* cells by both wildtype Drosha and N-terminal truncated Drosha (390-1374). (G) Restoration of *mir-144/451* processing in *Drosha/DGCR8-dKO* cells (from Shuo Gu lab) by wildtype or N-terminal truncations of Drosha (391-1374) and DGCR8[146-773]. (H) DGCR8 can co-IP with functionally active SAFB2[512-726], and this interaction was substantially strengthened in *Drosha-KO* cells.

It was reported that SAFB2 and Drosha interact, with the minimal regions corresponding to SAFB2[512-726] and Drosha[1-390] (Hutter et al. 2020) (**Figure 4A**). However, our co-IP tests did not detect substantial interactions between functional SAFB2[512-726] with either full-length or N-terminally truncated Drosha, in either wildtype or Microprocessor-knockout cells (**Figure 4D** and **Supplementary Figure 5A**). To further test whether the interaction of Drosha and SAFB2 is required for cluster assistance. To test this, we performed rescue assays for suboptimal miRNA processing in *Drosha-KO* cells, using full-length or N-terminally truncated Drosha lacking the reported SAFB2 binding region (**Figure 4E**). Interestingly, Drosha[391-1374] restored the processing of optimal *mir-144* and suboptimal *mir-451* in *Drosha-KO* cells, to levels equivalent to full-length Drosha (**Figure 4F**). Accordingly, neither the N-terminal region of Drosha nor its potential interaction with SAFB2 is essential for suboptimal miRNA processing.

We performed more stringent tests by expressing N-terminal truncation variants of Drosha and DGCR8 in *Drosha/DGCR8* double knockout (dKO) cells. These cells cannot mature endogenous nor exogenous canonical miRNAs, and remain fully incompetent even when singly expressing Drosha or DGCR8 (**Figure 4G**). Only upon co-expression of Drosha and DGCR8 do we observe normal maturation of ectopically expressed *mir-144/451* and *mir-375*, along with endogenous let-7a and miR-16 (**Figure 4G**). Co-expression of either N-terminal deletion variant of Drosha and DGCR8 (DroshaΔN or DGCR8ΔN) along with the reciprocal full-length partner provided near-normal rescue. Remarkably, co-expression of DroshaΔN and DGCR8ΔN also substantially rescued *Drosha/DGCR8-dKO* cells, including for processing of suboptimal *mir-451* (**Figure 4G**). While the efficiency of miR-451 processing was slightly lower than with rescue by full-length Microprocessor, we find that other canonical miRNAs, both exogenous and endogenous, were similarly compromised (**Figure 4G**). Therefore, dual N-terminally truncated Microprocessor does not function quite as well as wildtype, but the combined loss of their ERH and SAFB2-interacting domains does not specifically compromise cluster assistance.

Adding further complexity to the network of potential interactions amongst Microprocessor cofactors, ERH was reported to associate with the C-terminal RG/G-rich regions of SAFB1/2 (Drakouli et al. 2017). However, steady-state interactions between ERH and SAFB2[512-726] were weak at best, in either wildtype or Microprocessor mutant cells (**Supplementary Figure 5B**). Since SAFB2[512-726] is competent for cluster assistance, but lacks the RG/G-rich region, the potential interaction between ERH and SAFB2 seems not to be required for cluster assistance.

If SAFB2 does not mediate cluster assistance via the reported interactions with Drosha or ERH, how might it interface with Microprocessor? Unexpectedly, we found that SAFB2 could co-IP DGCR8 modestly, but that this interaction was strongly enhanced in *Drosha-KO* cells (**Figure 4H**). Although SAFB2 is not required for DGCR8 dimerization, nor for interaction of Drosha and DGCR8 (**Supplementary Figure 5C**), this still provides a possible means for SAFB2 to recruit Microprocessor at suboptimal miRNAs.

### Separable functions for ERH and SAFB2 during miRNA cluster assistance

We proposed a Microprocessor “Recruit-Transfer” model for cluster assistance, involving the following. Step 1: recruitment of Microprocessor and cleavage of the optimal miRNA hairpin, Step 2: release and transport of Microprocessor to the suboptimal miRNA hairpin, and Step 3: recognition and processing of suboptimal miRNA hairpin by Microprocessor (Shang et al. 2020). This framework is agnostic to the roles of cofactors such ERH and SAFB2, which could either assist one or more steps in this model, or conceivably play orthogonal roles (**Supplementary Figure 1**). We used tethering assays in our panel of mutant cells to dissect settings that do, or do not, require these cofactors (**Figure 5A**).

**Figure 5.**
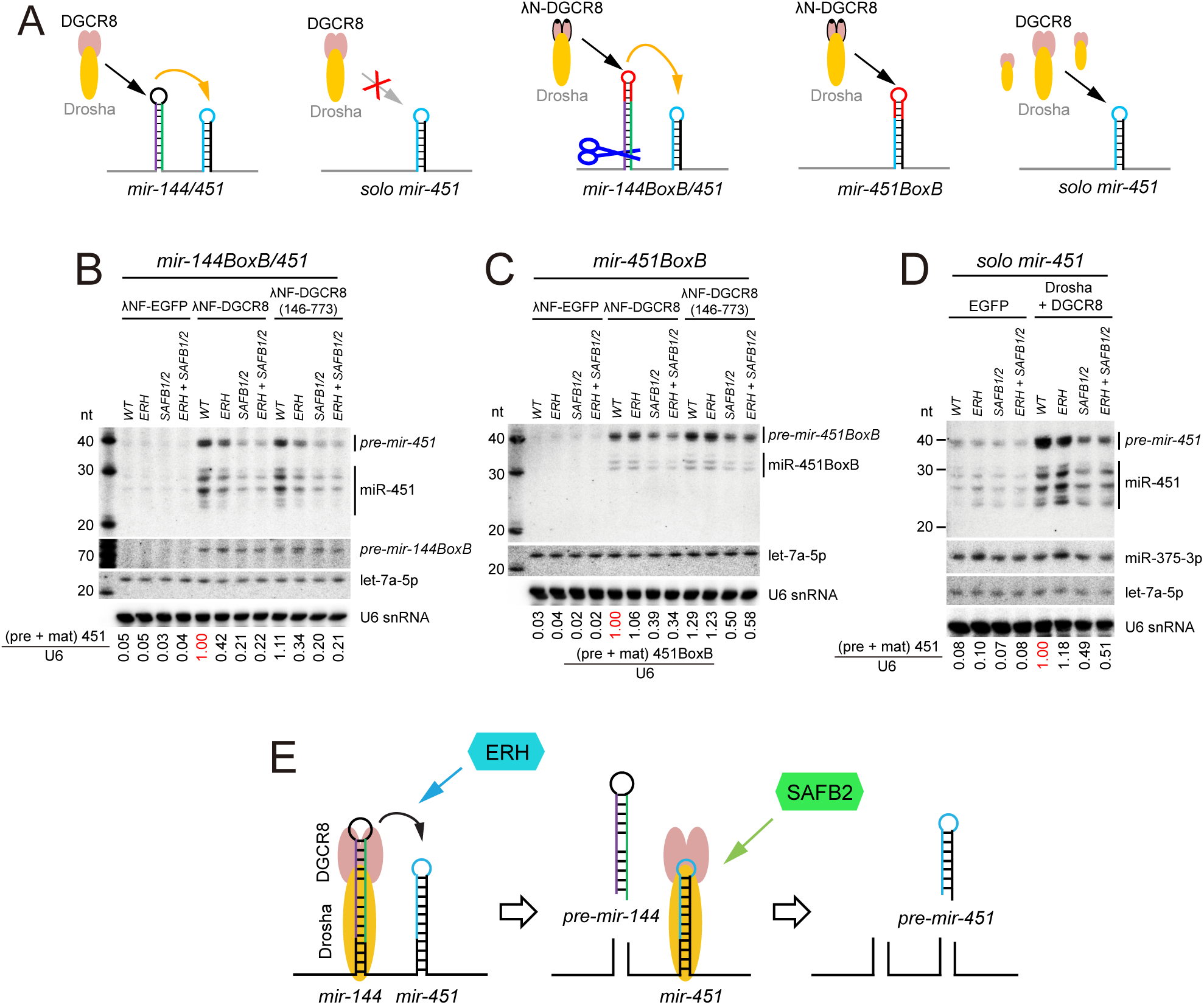
ERH and SAFB2 operate at distinct steps of cluster assistance. (A) Schematics of variant *mir-451* expression constructs, some of which are modified to contain a BoxB element that recruits λN-DGCR8 and thus Microprocessor. (B) Direct tethering of either full length λN-DGCR8 or N-terminal truncated λN-DGCR8[146-773] to the neighboring miRNA hairpin in the cluster can efficiently enhance suboptimal *mir-451* processing in WT cells. In this setting, both ERH and SAFB2 are needed for functional cluster assistance. (C) Tethering λN-DGCR8 to the solo *mir-451BoxB* hairpin also promotes its processing, and now bypasses the requirement for ERH. However, SAFB factors are still required in this context. Tethered λN-DGCR8[146-773] showed similar genetic dependencies for solo *mir-451* processing, further demonstrating that ERH is dispensable for recognition and processing of suboptimal miRNA hairpin. (D) Overexpression of Microprocessor can enhance *mir-451* processing in the absence of neighboring optimal *mir-144*. In this context, SAFB factors but not ERH are required for enhanced biogenesis of solo *mir-451*. This confirms that SAFB factors function after Microprocessor finds the suboptimal miRNA hairpin. (E) Model for sequential roles of ERH and SAFB2 during miRNA cluster assistance. In Step 1, Microprocessor is recruited by the optimal miRNA hairpin for its processing, which does not involve the cofactors. In Step 2, the transfer of the Microprocessor to the suboptimal miRNA hairpin is assisted by ERH. In Step 3, the suboptimal miRNA hairpin is recognized and processed by Microprocessor with the help of SAFB2, potentially via interaction with DGCR8.

As shown in **Figure 1B**, the biogenesis of *pre-mir-144* does not require ERH or SAFB2, reflecting that these factors do not impact the first step of Microprocessor recruitment and cleavage. We took advantage of this by directly tethering Microprocessor, via λN-DGCR8 to *mir-144* using a BoxB element (**Figure 5A**). Nuclear cleavage of *mir-144BoxB* hairpin was similar in wildtype and mutant cells (although this extension blocked subsequent processing by Dicer), while suboptimal *mir-451* was efficiently processed only in wildtype cells (**Figure 5B**). Curiously, recruiting Microprocessor with N-terminally truncated λN-DGCR8[146-773] still yielded *pre-mir-144BoxB* hairpin regardless of the cofactors, but only strongly enhanced *mir-451* in the presence of both ERH and SAFB2 (**Figure 5B**). This provided evidence that ERH and SAFB2 enhance suboptimal miRNA processing, after Microprocessor recruitment and cleavage of the optimal miRNA hairpin, and in a manner independent of ERH and DGCR8 interaction.

We next separated steps 2 and 3 in the “Recruit-Transfer” model, by tethering Microprocessor to solo *mir-451BoxB* (**Figure 5A**); i.e. without neighboring *mir-144* (Shang et al. 2020). With this substrate, we found that SAFB2, but not ERH, was required (**Figure 5C**). Moreover, direct recruitment of λN-DGCR8[146-773] to *mir-451* efficiently enhanced its processing, further indicating that interaction of ERH and DGCR8 can be bypassed during cluster assistance (**Figure 5C**).

Finally, we coexpressed solo *mir-451* with Microprocessor. In wildtype cells, this condition can dramatically enhance biogenesis of a suboptimal solo miRNA (Shang et al. 2020). In the presence of ectopic expression of Microprocessor in cofactor mutant cells, the biogenesis of miR-451 could bypass ERH, but not SAFB2 (**Figure 5D**). We conclude that ERH is not involved in the recognition and processing of suboptimal miRNA hairpin in step 3, but instead mediates the movement of Microprocessor between optimal and suboptimal hairpins in step 2. On the other hand, we infer that SAFB2 mainly promotes recognition and/or cleavage of the suboptimal miRNA hairpin in step 3, although it may also play a role during Microprocessor transport. Altogether, these tests clearly reveal separable and likely sequential roles for ERH and SAFB2 during miRNA cluster assistance (**Figure 5E**).

### Global effects of ERH and SAFB2 on miRNA expression

Previous studies of global effects of ERH and SAFB1/2 on endogenous miRNAs, involved manipulations of different cell lines (HEK293T, K562, hMSC and Ramos cells) (Fang and Bartel 2020; Hutter et al. 2020), which complicates their meta-analysis (Kwon et al. 2020). Moreover, mutant combinations of ERH with SAFB1/2 have not yet been assessed. To enable direct comparisons, we generated biologically replicate small RNA datasets across our panel of HEK293T mutant settings, namely *ERH* single depletion, *SAFB1/2* double depletion, and *ERH+SAFB1/2* triple depletion. Importantly, to be able to assess biogenesis effects on newly-synthesized miRNAs, we used SLAM-seq (Herzog et al. 2017; Reichholf et al. 2019) with 6 hr labeling. As we sequenced these data deeply (∼60M reads per library), we were able to broadly assess both steady-state and nascent populations of miRNAs (11-15M reads per library). We provide a number of processing statistics and quality control analyses of these datasets in **Supplementary Figure 6A-G**. Importantly, the replicates for each genotype were highly correlated for the abundance of individual miRNAs (**Supplementary Figure 6H**). Detailed statistics of the libraries are provided in **Supplementary Table 1**.

As a preliminary check of our data, we performed principal component analysis (PCA) and found that the replicates were well-correlated and the genotypes substantially distinct from each other (**Figure 6A**). The *SAFB1/2* and *ERH+SAFB1/2* mutant settings were far more separated from wildtype along PC1 than *ERH* mutants, but the two genotypes with *SAFB1/2* loss separated from each other on the PC2. We then assessed the behavior of miRNAs previously reported to be sensitive to ERH and/or SAFB function from meta-analysis of data collected from different cell types (Kwon et al. 2020). Our data largely agree with published data, although we generally find larger effect sizes for dependence on SAFB1/2 (**Supplementary Figure 7**). We also note that the triple *ERH+SAFB1/2* mutant condition appeared to have milder changes for several miRNAs than *SAFB1/2* mutants. While this nominally represents a rescue, given that ERH and SAFB1/2 have numerous functional roles beyond miRNA cluster assistance, this might also reflect a combination of indirect consequences in these mutant cells. For example, our data also recapitulate that a few miRNAs are upregulated in *ERH* and/or *SAFB1/2* mutants (Kwon et al. 2020) (**Supplementary Figure 7**), which we tentatively assign as indirect effects.

**Figure 6.**
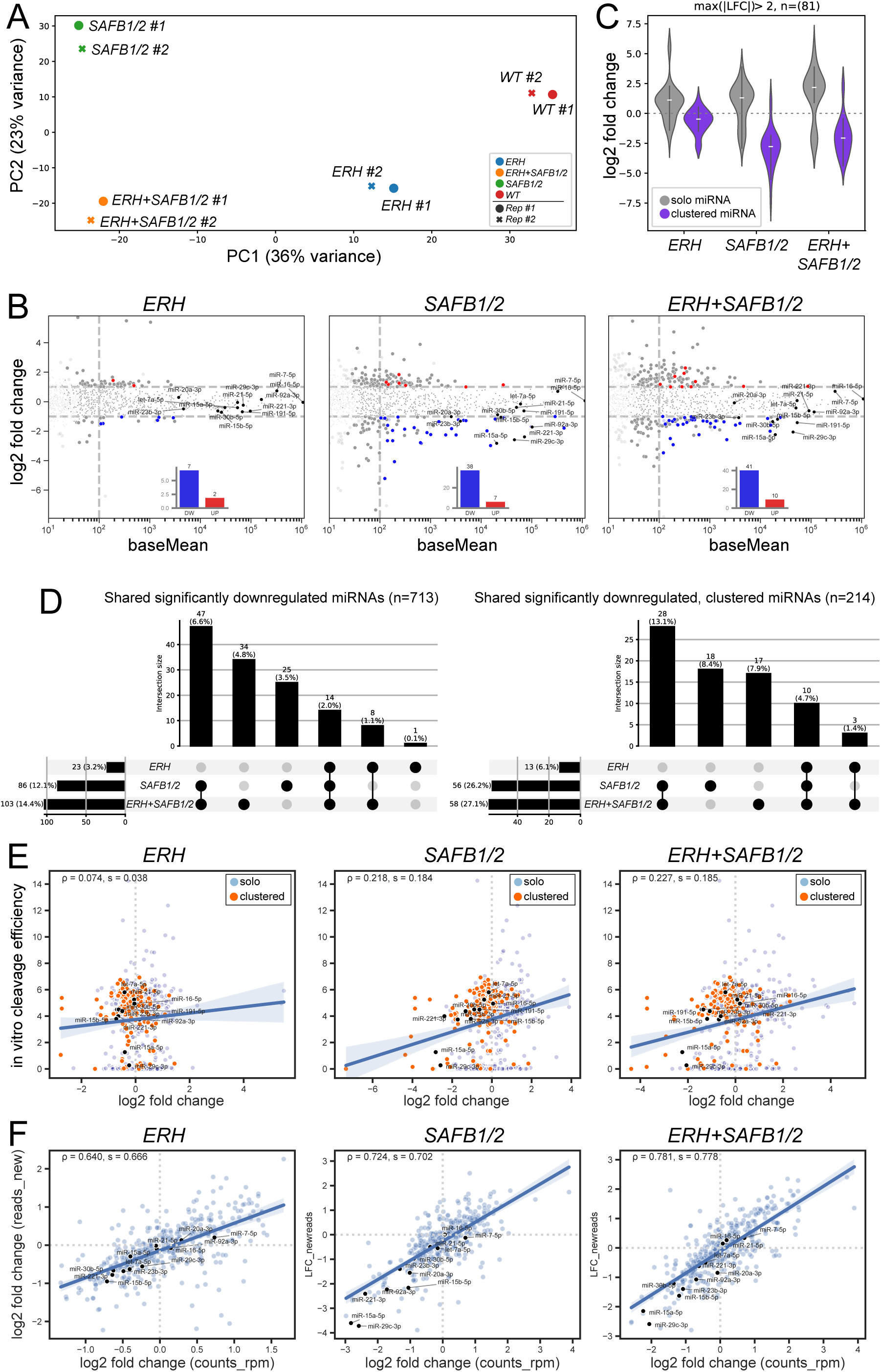
Global effects of ERH and SAFB1/2 on endogenous miRNAs. (A) PCA analysis of RPM-normalized read counts across the replicate small RNA libraries from four genotypes of HEK293T cells. (B) MA plots showing all tested (loci with >50 raw reads, n=713) and all clustered significantly downregulated (Adjusted P-Value p_adj_<0.05, log_2_ fold change LFC<-1; blue) and upregulated (p_adj_<0.05, LFC>1; red) miRNAs with average normalized counts (baseMean) ≥ 100. MiRNAs that were tested by Northern blotting in this study are annotated and highlighted using black color, miRNAs that were filtered in the final dataset (see Methods) are plotted transparently, miRNAs with baseMean < 10 are omitted for clarity. Clustered miRNAs are enriched amongst the downregulated miRNAs, especially upon loss of SAFB factors. (C) Violin plots of miRNAs whose absolute value changed by >2 fold (|LFC|>1) in the respective mutants, highlighting the preferential downregulation of clustered miRNAs. (D) UpSet plots showing the overlaps of differentially expressed miRNAs amongst the genetic manipulations. Loss of SAFB1/2 has the most substantial impact, but many miRNAs are codependent on ERH and SAFB1/2. (E) Correlations of ERH and/or SAFB1/2 dependencies with efficiency of *in vitro* cleavage by recombinant Microprocessor+SRSF3. We analyzed 487 miRNAs that had both *in vitro* cleavage data and sufficient reads in the deep sequencing data. The positive correlation between these datasets, especially with SAFB1/2 dependency, indicates that these Microprocessor cofactors facilitate suboptimal Microprocessor substrates. Pearson and Spearman correlation coefficients are denoted by ρ and s, respectively. (F) Comparison of steady state and nascent (<6 hours) miRNA expression in mutants of Microprocessor cofactors with sufficient nascent reads (440 loci). Positive correlation between these populations supports that these cofactors have direct impacts on miRNA biogenesis. For reference, miRNAs whose expression was validated by Northern blotting in this study are labeled in black.

Since our data are deeper than previous assessments of Microprocessor cofactor depletions (Fang and Bartel 2020; Hutter et al. 2020; Kwon et al. 2020), we were able to assign more differentially expressed miRNAs. MA plots (**Figure 6B**) and volcano plots (**Supplementary Figure 8**) summarize that loss of SAFB1/2, with or without simultaneous ERH depletion, affected many more miRNAs than did ERH depletion alone. We selected several classes of miRNA expression changes for validation. Interestingly, for the most part, we observed similar changes in the corresponding miRNA* strands, consistent with these reflecting altered processing (**Supplementary Figure 9**). The endogenous miRNAs we examined by Northern blotting include ones that were preferentially downregulated in SAFB1/2 mutants (miR-15a, miR-29c, miR-221), ones that were dependent on both ERH and SAFB1/2 (miR-15b, miR-23b, miR-30b, miR-191), one that was increased in mutants (miR-7), and a control that was relatively unchanged in mutants (miR-16). These experimental tests validated the changes measured from deep sequencing (**Supplementary Figure 9**). While altered miRNAs spread in both directions, we note that clustered miRNAs were prominently enriched amongst the downregulated miRNAs. This is more clearly visualized in violin plots of the respective mutant genotypes, where we stratified miRNA accumulation by cluster status (**Figure 6C**). Overall, these data are consistent with the notion that these Microprocessor cofactors play specialized roles to support miRNA cluster assistance.

It was earlier concluded that most suboptimal miRNAs are co-dependent on ERH and SAFB1/2 (Kwon et al. 2020). We used UpSet plots to visualize the genetic dependencies of miRNAs, and these highlight that the majority of downregulated miRNAs are dependent on SAFB1/2, regardless of whether ERH was present or not, and that additional miRNAs were SAFB1/2 alone (**Figure 6D** and **Supplementary Figure 10**). On the other hand, only a small minority of downregulated miRNAs were changed only in *ERH* or *ERH+SAFB1/2* depletion conditions. Although a caveat is that these are strong loss-of-function but not null conditions, due to cell lethality of all of these factors in HEK293T (**Figure 1A**), these data support the notion that SAFB1/2 play larger roles for cluster assistance than ERH. Moreover, they tentatively suggest that SAFB factors impact miRNA biogenesis downstream of ERH, in line with our other data (**Figures 1** and **5**).

### Profiling from ERH and/or SAFB1/2 mutant conditions reveals suboptimal miRNAs

As noted, these mutant profilings do not guarantee that miRNA expression is altered due to defective cluster assistance. For example, miR-1226-3p was reproducibly downregulated ∼20% in *ERH* and 50% in *ERH+SAFB1/2* mutants (**Supplementary Table 1**). Since this is the mature strand of a mirtron, which uses splicing to bypass Microprocessor for nuclear biogenesis (Berezikov et al. 2007; Westholm and Lai 2011), this change should not be due to cluster assistance *per se*. Thus far, the only direct assay of suboptimality of nuclear miRNA processing is to show that the biogenesis of a given hairpin can be increased when in proximity to an optimal neighbor. This readout shows that even some “solo” miRNAs are suboptimal Microprocessor substrates (Shang and Lai 2023). However, such assays are not yet scalable.

To gain insights as to whether dependence on Microprocessor cofactors reflects suboptimality as Microprocessor substrates, we took advantage of high throughput *in vitro* Microprocessor cleavage data (Kim et al. 2021). Such cleavage data have a limitation in that the reaction conditions of recombinant proteins and synthetic RNAs are not expected to be the same as in cells (Shang et al. 2020). We observed a mildly positive correlation of ERH dependency with Microprocessor cleavage efficiency (**Figure 6E**, left panel). However, we observed stronger correlations upon comparing *in vitro* cleavage efficiency with genetic dependencies on SAFB1/2 or ERH+SAFB1/2 (**Figure 6E**, middle and right panels). This provides evidence that our profiling data helped to classify functionally suboptimal miRNAs that are enhanced by Microprocessor cofactors (**Supplementary Table 2**).

We summarize examples of ERH and/or SAFB1/2-dependent miRNAs bearing suboptimal features in **Supplementary Figure 11**. We selected *mir-92a* for validation, as a locus strongly dependent on SAFB1/2, but largely independent of ERH. Its behavior fits with our structure-function data (**Figure 2**), since *pri-mir-92a-1* exhibits a clearly suboptimal lower stem structure. We tested its accumulation across our mutant cell lines using Northern blotting, and confirmed strong reduction of mature miR-92a in *SAFB1/2*-depleted, but not *ERH*-depleted cells (**Figure 7**). This provides further evidence of the separable functions of these Microprocessor cofactors. In fact, for reasons unknown, *ERH+SAFB1/2* depleted cells accumulate higher levels of miR-92a than *SAFB1/2*-depleted cells. The basis of this potentially antagonistic effect remains to be clarified, but there might be further complexities in cluster biogenesis. For example, we have documented cases of Microprocessor competition (instead of collaboration) within clusters (Shang et al. 2020; Shang and Lai 2023).

**Figure 7.**
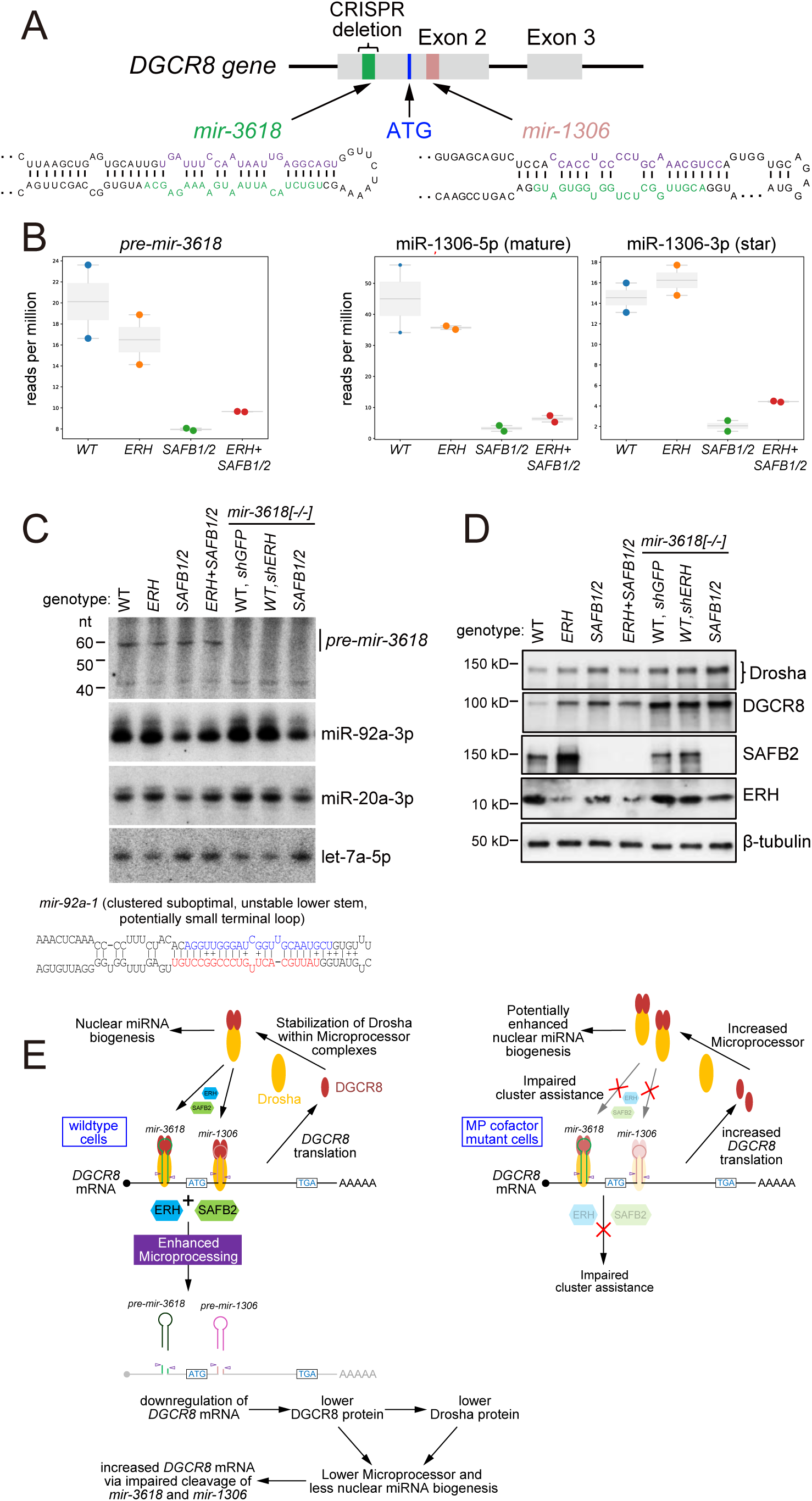
Impact of miRNA cluster assistance on Microprocessor homeostasis. (A) The DGCR8 transcript contains two nearby hairpins that are cleaved by Drosha. These are formally classified as miRNAs but inefficiently generate mature miRNAs, suggesting that their major function may be to regulate DGCR8 levels during Microprocessor homeostasis. *mir-3618* resembles a canonical hairpin but its pre-miRNA is reported to be nuclearly retained, while *mir-1306* is a clearly suboptimal hairpin with poor lower stem pairing. (B) Expression of small RNAs and pre-miRNAs from *ERH* and/or *SAFB1/2* mutant libraries. Mature miR-3618 and *pre-mir-1306* were negligibly expressed (<0.25RPM in all cases, usually less) and were thus not reliably quantified. (C) Northern blotting in wildtype and mutant settings for Microprocessor cofactors, as well as in deletion backgrounds for *mir-3618*. These tests validate specificity of *pre-mir-3618* signals and confirm absence of mature miR-3618 (see whole blots in **Supplementary Figure 13**). These blots also show that miR-92a is selectively and substantially decreased in any condition with depletion of SAFB1/2; *pri-mir-92a* exhibits a suboptimal lower stem. (D) Western blotting of Microprocessor components and its factors. DGCR8 and Drosha proteins are increased upon depletion of ERH and SAFB1/2. Deletion of *mir-3618* increases DGCR8 and Drosha proteins even more robustly, but loss of this *DGCR8* hairpin blunts the effects of ERH and SAFB1/2 loss. (E) Model for complex feedback regulatory interactions that maintain Microprocessor (MP) homeostasis, including the impact of ERH/SAFB1/2-mediated cluster assistance on *DGCR8* mRNA cleavage by Microprocessor.

Another major reason that miRNAs might change in steady state profilings is altered stability, as opposed to biogenesis defects. To address these, we analyzed miRNAs using the SLAM-seq labeling information aspect of our data. We used a statistical modeling framework for small RNA SLAM-seq to quantify the proportion of reads carrying 4SU-induced T>C conversions, thereby providing an estimate of the fraction of nascent versus pre-existing molecules (Herzog et al. 2017; Reichholf et al. 2019). This allowed statistical inference of the portion of unlabeled reads that were exposed to 4SU and thus distinguished between newly synthesized and pre-existing miRNAs. We recovered a substantial population of newly-synthesized miRNAs, which enabled broad assessment of their expression changes. As a quality control, we observed that miRNA* strands showed higher labeling fractions than their corresponding mature strands at the 6 hr SLAM-seq timepoint (**Supplementary Figure 12**), consistent with their overall faster turnover following maturation of Argonaute complexes into the mature single-stranded form (Reichholf et al. 2019).

Having verified the quality of our SLAM-seq data, we compared miRNA expression in the newly-synthesized and steady-state pools. These analyses revealed that differential miRNA expression was strongly correlated between all three mutant genotypes (*ERH*, *SAFB1/2* and *ERH+SAFB1/2*), compared to control HEK293T cells (**Figure 6F**). Accordingly, these data confirm a dominant impact of depleting cluster assistance factors on miRNA biogenesis. Altogether, our datasets expand our knowledge of ERH and/or SAFB1/2-dependent miRNAs, including suboptimal loci.

### Impact of suboptimal miRNA biogenesis on Microprocessor homeostasis

In our final assays, we sought to characterize a regulatory output that is modulated by cluster assistance. Several clusters bearing suboptimal miRNAs are associated with phenotypically relevant biology (e.g. *mir-144/451*, *mir-15a/16* and *mir-17∼92a*). In such contexts, ERH and SAFB1/2 levels might impact the repression of mRNA targets of suboptimal miRNAs. However, there are clear challenges to connect how changes in miRNA levels might be relayed to changes in their targets, given (1) the distributed nature of miRNA target networks and (2) the fact that ERH and SAFB1/2 have multiple roles outside of cluster assistance.

We therefore considered whether Drosha-mediated mRNA cleavage might be a favorable context to document *in vivo* regulatory impacts of cluster assistance. The first and most robust example of this is embedded within reciprocal cross-regulatory interactions between the two Microprocessor components. Microprocessor (via the ribonuclease Drosha), cleaves hairpins within the *DGCR8* transcript to repress DGCR8 protein levels, while DGCR8 stabilizes Drosha protein within the Microprocessor complex. This homeostatic circuit operates in both vertebrates (Han et al. 2009; Triboulet et al. 2009) and intact *Drosophila* (Kadener et al. 2009; Smibert et al. 2011). Although the layout of two *DGCR8* hairpins in mammals and flies is conserved, the miRNAs themselves are not, suggesting that cleavage of *DGCR8* mRNA is more fundamental than function of the encoded miRNAs.

In vertebrate *DGCR8*, *mir-3618* is located within the 5’ UTR while *mir-1306* resides in the N-terminal encoding region (**Figure 7A**). When first studied, these hairpins were confirmed as Drosha substrates, but already noticed to be less robustly processed than canonical pri-miRNA substrates (Han et al. 2009; Triboulet et al. 2009). In fact, *mir-1306* was noted to be less amenable to Drosha cleavage than *mir-3618*, even though currently only *mir-1306* (but not *mir-3618*) is included in curated mirGeneDB annotations (Clarke et al. 2025). The structure of *mir-1306* is particularly suboptimal, with poor lower-stem structure, modestly paired duplex stem, and a small terminal loop (**Figure 7A**).

Although our profiling data was aimed at characterizing small RNAs, we noticed that they captured a minor subset of pre-miRNA reads (**Supplementary Figure 6E**). Although mature miR-3618 reads were negligible, in line with previous reports that *pre-mir-3618* is predominantly retained in the nucleus (Han et al. 2009), there were in fact substantial *pre-mir-3618* reads in our sequencing data. These revealed a mild reduction of *pre-mir-3618* in *ERH* mutants, but a prominent >2-fold reduction in SAFB1/2 and ERH+SAFB1/2 mutant cells (**Figure 7B**). With *mir-1306*, its pre-miRNA reads were ∼100 fold lower than with *mir-3618*, and could not be assessed. However, we clearly observed that miR-1306 requires SAFB-mediated cluster assistance, since both its mature and passenger strand reads were reduced 5-10 fold in *SAFB1/2* and *ERH+SAFB1/2* mutant cells (**Figure 7B**). Although effects of ERH on miR-1306 were more subtle than with SAFB1/2, we note that miR-1306 was the most significantly reduced miRNA in prior analysis of *ERH*-knockdown cells (Kwon et al. 2020); however, its fold changes were similarly modest in that study.

We could not detect mature miR-3618 or miR-1306 by Northern blotting, even upon sensitizing with ectopically expressed constructs (**Supplementary Figure 13**). However, we detected a band of the expected size for the *pre-mir-3618* hairpin (Han et al. 2009), along with a shorter band. To establish their provenance, we used CRISPR/Cas9 to delete *pre-mir-3618* from wt and *SAFB1/2* mutant cell lines (**Supplementary Figure 2**). This eliminated the ∼58nt band but did not affect the ∼42nt band, validating the former as *pre-mir-3618* and the latter as a cross-reacting species (**Figure 7C**).

We then examined effects on the accumulation of Microprocessor proteins. Our hypothesis was that enhanced cluster assistance would increase *DGCR8* transcript cleavage, thus compromising accumulation of DGCR8 and Drosha proteins. Indeed, ERH and SAFB1/2 mutant conditions upregulated both Microprocessor proteins, although the effects were greater for DGCR8 (**Figure 7D**), suggesting that it is the proximal target of cluster assistance. However, it is also possible that this reflects processing inhibition of a miRNA-like hairpin in the Drosha mRNA (Lee et al. 2017; Mechtler et al. 2017).

To more directly test if these changes were due to cluster assistance *per se*, we used our *mir-3618* deletion cells. Here, we hypothesize that loss of the optimal miRNA partner should (1) compromise repression of DGCR8 protein by Microprocessor, and (2) blunt the regulatory effect of the remaining suboptimal *mir-1306* hairpin. This is precisely what we observe: DGCR8 and Drosha proteins were misexpressed in all *mir-3618* deletion genotypes, while the effects of ERH and SAFB1/2 were reduced in the absence of *mir-3618*, compared to cells that carry this hairpin (**Figure 7D**). Overall, we conclude that one of the most overt utilizations of miRNA cluster assistance via Microprocessor cofactors is actually to mediate cross-regulatory interactions between Drosha and DGCR8 during Microprocessor homeostasis (**Figure 7E**).

## Discussion

### Complex and separable roles for ERH and SAFB2 in miRNA cluster assistance

Since the discovery of cluster assistance for suboptimal miRNA biogenesis (Fang and Bartel 2020; Hutter et al. 2020; Shang et al. 2020), further mechanistic insights have been limited. A major reason for this is the complexity of this process, which involves the Microprocessor (Drosha and DGCR8), at least two cofactors (ERH and SAFB2), at least two miRNA hairpins, and dynamic interactions amongst all of these. The proposed working models are all based on interactions of these cofactors with Microprocessor components, including ERH-DGCR8 (Kavanaugh et al. 2015; Fang and Bartel 2020; Kwon et al. 2020), and SAFB2-Drosha (Hutter et al. 2020) interactions, along with ERH-ERH dimerization (Arai et al. 2005; Wan et al. 2005; Kwon et al. 2020), SAFB2 dimerization (Hutter et al. 2020; Korn et al. 2023) and ERH-SAFB2 interaction (Drakouli et al. 2017). Accordingly, many plausible models of miRNA cluster assistance involve higher order complexes of Microprocessor (Shang et al. 2023; Kim et al. 2025). However, the various models cannot be distinguished from the reported data.

While we can recapitulate some of these interactions, our data indicated that many proposed interactions are weak or not stably detected. More importantly, we find that several reported protein-protein interactions are not essential for cluster assistance. It remains possible that these are transient interactions and/or play supporting roles in the process. For example, although we show that ERH-mediated dimerization of DGCR8 (Kwon et al. 2020) is not essential for cluster assistance, we also verify that dimerization of DGCR8 is indeed essential for canonical and suboptimal miRNA biogenesis. In fact, as three strategies were proposed for DGCR8 dimerization involving ERH (Kwon et al. 2020), heme binding (Garg et al. 2024) and the RHED domain (Faller et al. 2007), there are evidently fail-safe strategies to ensure this state. Nevertheless, our data that certain protein-protein interactions can be bypassed during effective cluster assistance help to distinguish correlative attributes from causal ones. Importantly, we were able to delineate a series of conditional requirements for Microprocessor cofactors, and showed that these differ between ERH and SAFB2. Our data support the notion that ERH and SAFB2 actually play separable roles in a sequence of events that underlie Microprocessor transfer during miRNA cluster assistance, with ERH playing a role in Microprocessor transfer between hairpins, and SAFB2 critical for stable recognition and/or processing of the suboptimal hairpin *per se*.

During review/revision of our study, related works on miRNA cluster assistance were published (Jang et al. 2025) or preprinted (Aschenwald et al. 2025). While it was previously reported that ERH mediates cluster assistance by binding the N-terminus of DGCR8 (Kwon et al. 2020), our study and work from the Herzog lab indicate that loss of the ERH-binding region of DGCR8 is compatible with cluster assistance (Aschenwald et al. 2025). In addition, work from the Kim lab concluded that SAFB2 operates during transfer of Microprocessor between hairpins (Jang et al. 2025), while our study support that SAFB2 is intrinsically required at the suboptimal miRNA hairpin for its processing (**Figure 5E**). Tests from the Herzog lab are consistent with the latter interpretation (Aschenwald et al. 2025).

In our small RNA sequencing data, we find certain “solo” miRNAs are preferentially dependent on ERH, and not on SAFB2 (**Supplementary Figure 11**). This may be taken to reflect roles of ERH in miRNA biogenesis beyond cluster assistance, as proposed in concurrent studies (Aschenwald et al. 2025; Jang et al. 2025). Currently, our tests do not distinguish against the possibility of indirect consequences on miRNA upon silencing ERH. However, our tests confirm that solo, suboptimal miRNAs do exist, and that their maturation can be enhanced by ectopic Microprocessor (Shang and Lai 2023), or by recruitment via ERH (**Figure 3I-J**). Of note, a recent study showed that SAFB can recognize an RNA hairpin in the *Nfib* transcript, thereby recruiting Drosha to cleave and downregulate *Nfib* in neural stem cells (Forcella et al. 2024). Thus, SAFB might also have role in hairpin processing beyond miRNA cluster assistance. Finally, we also find that some miRNAs are elevated upon depletion of ERH and/or SAFB1/2, such as miR-7 (**Supplementary Figure 9C**). Whether this is indirect, or reflects additional complexity of Microprocessor cofactors on miRNA biogenesis, deserves further study.

### Co-option of multiple regulatory factors for regulation of nuclear miRNA biogenesis

Both ERH and SAFB1/2 are multifunctional factors that are involved in numerous other regulatory processes. Moreover, some of their core features are preserved in species that lack miRNAs. For example, ERH dimerizes in *S. pombe*, where it assembles a complex with the YTH protein Mmi1 (Hazra et al. 2020). Similarly, mammalian ERH dimerizes and assembles with DGCR8 (Kwon et al. 2020). It is therefore useful to provide some perspective on other processes that utilize these factors.

ERH is well-conserved across eukaryotes, and ranges from 100-113 aa (104 aa in human), nearly qualifying as a “microprotein”. It was originally identified using *Drosophila* genetics, as an enhancer of the *rudimentary* wing defect caused by a defect in pyrimidine biosynthesis (Wojcik et al. 1994), and later found in mammals, plants and fungi (Kozlowski 2023). ERH resides in both cytoplasm and nucleus (Pogge von Strandmann et al. 2001), and can be selectively recruited to subcompartments such as nuclear speckles (Banko et al. 2013). Accordingly, ERH has been linked to a diverse array of regulatory processes including transcriptional regulation, splicing control, mRNA decay (Kozlowski 2023). Strikingly, a recent study reported that ERH was one of the most highly interconnected factors amongst RBP-centered complexes, extending far beyond previously reported SAFB1/2 and DGCR8 (Street et al. 2024). Thus, ERH qualifies as a genuine regulatory hub.

Intriguingly, the *C. elegans* piRNA (21U RNA) biogenesis pathway, which bears little relationship to other metazoan piRNA pathways except for the fact that the resultant small RNAs assemble with a Piwi-class effector protein, has co-opted an ERH homolog. In particular, ERH-2 homodimerizes as a constituent of two regulatory complexes with overlapping content (Cordeiro Rodrigues et al. 2019; Perez-Borrajero et al. 2021). While ERH-2 binds PID-3 in the common PETISCO complex, ERH-2 is the adaptor for two additional partners (PID-1 and TOST-1) that bind in a mutually exclusive fashion through the same interface. PID-1 confers activity in 21U biogenesis, while TOST-1 is important for embryonic viability that is independent of piRNAs. These findings highlight how ERH proteins have been independently co-opted for regulation of small RNA biogenesis, and how these miniature proteins surprisingly have sufficient “real estate” to coordinate multiple distinct protein complexes.

SAFB1 (also known as SAFB) and SAFB2 are highly similar paralogs that are arranged head-to-head on human chromosome 19 (Renz and Fackelmayer 1996; Townson et al. 2003). Not only can both factors homodimerize, they are members of a select group of proteins with capacity for independent recognition of DNA and RNA, accomplished via their SAP and RRM domains, respectively (Korn et al. 2023). As integral components of the nuclear matrix in vertebrate cells, both SAFB factors can bind scaffold/matrix attachment region (S/MAR) DNA. However, they also play significant roles in RNA regulation, including splicing and binding of nascent RNA (Townson et al. 2003; Sergeant et al. 2007; Rivers et al. 2015; Cherney et al. 2023). They are collectively involved in cell cycle regulation, apoptosis, differentiation, the stress response, regulation of immune genes (Norman et al. 2016), as well as transposon control (Hong et al. 2024; Ilik et al. 2024).

Overall, it seems sensible that ERH and SAFB1/2 could be co-opted into the miRNA pathway, since it is already clear that these versatile factors integrate into many other pathways. We showed that they actually have separable roles during cluster assistance, which may also be logical if we imagine they were independently incorporated into this process. Thinking more broadly, we can speculate that even though the core of dedicated miRNA biogenesis factors was characterized long ago, there are still abundant opportunities for other proteins with other established functions to “moonlight” as part of the miRNA pathway.

### Broader impacts and evolutionary implications of cluster assistance

Cluster assistance facilitates the processing of suboptimal miRNAs not only in mammals, but also for miRNAs in *Drosophila* and some viruses (Haar et al. 2016; Truscott et al. 2016; Vilimova et al. 2021). Currently, only about a dozen suboptimal miRNAs whose biogenesis is enhanced by proximity to an optimal neighbor have been experimentally validated. As >1/3 of animal miRNAs reside in clusters, presumably more await characterization. Importantly, loss of ERH and SAFB1/2 can perturb miRNA expression independently of cluster assistance *per se*, and there may be additional strategies to enhance suboptimal miRNAs. Thus, direct manipulations of miRNA loci remain critical to document such regulation. A high-throughput strategy involving massive parallel screening would be critical to systematically reveal all the suboptimal miRNAs in different tissues and species.

The other intriguing open question is why cluster assistance is selected and kept for miRNA processing during evolution? One possibility is that young miRNA genes might be expected to be suboptimal, as *de novo* loci would not have been evolutionarily selected for optimal biogenesis features. Emergence within a cluster could facilitate the maturation of such a suboptimal miRNA (Mohammed et al. 2014; Shang et al. 2020), and permit evolutionary selection for beneficial targeting that could in turn promote enhanced biogenesis of the miRNA (Bartel and Chen 2004). This scenario may help explain young suboptimal miRNA hairpins that reside in miRNA clusters. But other suboptimal miRNA genes, like *mir-451* and *mir-15a*, are conserved across vertebrates, yet maintain their suboptimal features. Why haven’t these transitioned into optimal miRNA hairpins? One generic explanation is that this allows greater control over miRNA biogenesis and thus gene regulation. But a more specific rationale is that this enables hierarchical control of miRNA biogenesis. As we showed, suboptimal miRNAs are processed after their optimal partners (Shang et al. 2020; Shang and Lai 2023), and there might be settings in which tuning of cluster assistance mechanisms could enable selective expression of the optimal members of a miRNA operon. Finally, by virtue of its atypical Dicer-independent biogenesis strategy, *mir-451* may be locked into utilizing cluster assistance. In fact, both *mir-144* and *mir-451* are subject to both independent and intertwined layers of biogenesis regulation (Yoda et al. 2013; Kretov et al. 2020; Shang et al. 2022), which may be relevant for their functions during erythropoiesis.

It is likely that other factors and strategies for enhancement of nuclear miRNA biogenesis exist. For example, our data indicate that ERH does not require direct interaction with DGCR8 to support cluster assistance via Microprocessor transfer. It is conceivable that this involves one or more other factors documented amongst the extensive ERH interaction network (Street et al. 2024). In addition, we previously showed that the solo miRNA *mir-491* harbors a functionally suboptimal terminal loop that impedes Microprocessor recruitment (Shang and Lai 2023). While the biogenesis of *pri-mir-491* can be strongly enhanced by a neighboring optimal miRNA, this strategy of cluster assistance is not available in its endogenous genomic location. We speculate that *pri-mir-491* may recruit another unknown factor that facilitates recruitment and/or cleavage by Microprocessor. Indeed, there could be many miRNAs that utilize trans-acting factors, whose identification and functional delineation await characterization (Treiber et al. 2017; Nussbacher and Yeo 2018).

## Materials and Methods

### Plasmid constructs

Plasmids for expression of all the miRNAs or miRNA clusters used in this study were constructed by inserting amplified DNA fragments containing the miRNA precursors (200 bp∼1 kb) from genomic DNA of HEK293T cells between Bgl II and Xho I sites downstream of a CMV promoter. All the miRNA mutants were constructed using overlapping PCR method based on the wildtype miRNA constructs. The cDNA plasmids used in this study were constructed with HA, Flag and/or λN peptides on their N-terminal or C-terminal ends. The shRNA plasmids used in this study are constructed by directly inserting the annealed DNA oligos downstream of a U6 promoter (Shang et al. 2015). The CRISPR plasmids used in this study are constructed by directly inserting the annealed DNA oligos into the lentiCRISPRv2 vector with a GFP or puromycin resistant selection marker. All the details and oligonucleotide sequences used to clone these constructs are listed in **Supplementary Table 3**.

### Cell culture

Wildtype and mutated HEK293T cells were grown in DME-high glucose media containing 10% FBS, 1% non-essential amino acids, 1% sodium pyruvate, penicillin/streptomycin and 0.1% 2-mercaptoethanol. Mycoplasma contaminations were regularly tested for the cell lines.

### RNAi-mediated knockdown

To deplete remaining ERH or SAFB1 factors, we applied RNAi-mediated depletion to the relevant mutant HEK293T cells. We transfected 2 µg of shRNA plasmids targeting the endogenous genes or control shRNA plasmids using Lipofectamine 2000 (Thermo Fisher) in 6-well plates. 36 hrs post transfection, cells were split and seeded into new culture plates for further transfections, according to the specific experimental design.

### Generation of knockout HEK293T cell lines using CRISPR/Cas9

Plasmids for gRNA expression were constructed by inserting annealed DNA oligonucleotides containing guide RNA sequences targeting *ERH* or *SAFB1/2* genes into the BsmBI site of lentiCRISPRv2-GFP or lentiCRISPRv2-PuroR. 2 µg of these plasmids were transfected into HEK293T cells in 6-well plates using Lipofectamine 2000. After 16 hrs, puromycin (2 µg/mL) was added for cells transfected with lentiCRISPRv2-PuroR. After 48 hrs, the cells transfected with lentiCRISPRv2-GFP were collected for cell sorting to split the GFP-positive single cells into 96-well plates. After two weeks of culturing, the cells were used for genotyping and then Western blotting to identify the deletion or depletion colonies. Oligo sequences for sgRNA and genotyping are listed in **Supplementary Table 3**.

### Northern blotting

Co-transfection of miRNA cluster plasmids (2 µg/well for 6-well plate or 1 µg/well for 12-well plate) with control mir-375 plasmid (200 ng/well for 6-well plate or 100 ng/well for 12-well plate) were performed in HEK293T cells using Lipofectamine2000. Two or three days post-transfection, total RNA was prepared from the cells using Trizol reagent (Invitrogen). Equal amounts of total RNAs (10-15 µg) were mixed with 2x RNA loading dye, denatured at 95°C for 5 min, and then fractionated by electrophoresis on a 20% urea polyacrylamide gel in 0.5x TBE buffer, until the bromophenol blue dye run out of the gel. The gel was transferred to GeneScreen Plus nylon membrane (Perkin Elmer) at 300 mA for 1.5 hr, UV-crosslinked with 120,000 µJ of energy, baked at 80°C for 30 min. Blots were hybridized with γ-^32^P-labeled DNA probes against mature miRNA sequences in hybridization buffer (5x SSC, 7% SDS, 2x Denhardt’s solution) at 42°C overnight. We washed the membrane with Non-Stringent Wash Solution (3x SSC, 5% SDS, 10x Denhardt’s solution), followed by two washes with Stringent Wash Solution (1x SSC, 1% SDS). Each wash step is conducted at 42°C for 30 min. Membranes were sealed in plastic wrap and exposed to a phosphorimager cassette for 1∼3 days. To re-probe a blot, we washed the blot with 1% SDS at 80°C for 30 min, prior to hybridization with additional probe(s). All the probe sequences are listed in **Supplementary Table 3**.

### Co-immunoprecipitation assays

Wildtype or mutant HEK293T cells were grown in 6-well plates and transiently co-transfected with plasmids (1 µg for each) of Flag- or HA-tagged cDNAs of relevant genes. After 48 hrs, the cells were washed with phosphate-buffered saline (PBS) and harvested in lysis buffer containing 50 mM Tris-HCl [pH 7.4], 150 mM NaCl, 1 mM EDTA, 1% Triton X-100 and protease inhibitor cocktail. After rotation at 4°C for 20 min, each lysate was clarified by centrifugation at 14,000 × g at 4°C for 15 min. A total of 500 µl supernatant was mixed with 10 µl of anti-Flag M2 Agarose beads (Sigma) or anti-HA magnetic beads (Thermo Fisher) at room temperature for 2 hrs or at 4°C overnight. The beads were washed four times with Tris-buffered saline (TBS) and used for Western blotting analysis.

### Western blotting

Input/IP products or HEK293T cell lysates were separated on 4-20% Mini-PROTEAN TGX Precast Protein Gels (Bio-Rad) and then transferred to PVDF membranes. The blots were probed for 2 hrs at room temperature or overnight at 4°C with the relevant antibodies: rabbit polyclonal anti-SAFB1 (Thermo Fisher, Cat #A300-812A, 1:2000), rabbit polyclonal anti-SAFB2 (Thermo Fisher, Cat #A301-112A, 1:2000), or rabbit polyclonal anti-ERH (Thermo Fisher, Cat #PA5-21388, 1:2000), or rabbit monoclonal anti-DGCR8 (3F5) (Thermo Fisher, Cat #MA5-24860, 1:3000), rabbit monoclonal anti-Drosha (D28B1) (Cell Signaling Technology, Cat# 3364, 1:3000), or mouse monoclonal anti-β-tubulin (DSHB, 1:2000), followed by secondary antibody conjugated to horseradish peroxidase (1:5000). For Flag- or HA-tagged proteins, the blots were directly probed for 2 hrs at room temperature with HRP-conjugated mouse monoclonal anti-Flag (Sigma, Cat #A8592, 1:5000), or HRP-conjugated mouse monoclonal anti-HA (Thermo Fisher, Cat #26183-HRP, 1:5000). Signals were detected with Amersham ECL Prime Western Blotting Detection Reagent.

### SLAMseq (Thiol (<u>S</u>H)-<u>L</u>inked Alkylation for the <u>M</u>etabolic <u>seq</u>uencing of RNA) small RNA library construction

The wild-type, *ERH[+/-]*, *SAFB1[+/-]/SAFB2[-/-]* and *ERH[+/-]/SAFB1[+/-]/SAFB2[-/-]* HEK293T cells cultured in 6-well plates were first treated by shRNA transfections to further deplete endogenous ERH and SAFB1 as shown in Figure 1 for 48 hrs. Each genotype had two biological replicates. Then cells from each well were divided and half were replated for further culture, while the other half cells were collected for validation. The replated cells were cultured for another 24 hrs, and replaced with fresh medium containing 400 μM of 4SU to label the nascent cellular RNAs in dark for 6 hrs. Then cells were collected by Trizol for total RNA extraction, during which Reducing Agent (SLAM-seq Kinetics Kit-Anabolic kinetics module, Cat# M06124-2-0100) were added for maintaining the 4SU treated samples under constant reducing conditions.

Following addition of 4SU, all subsequent steps were protected from light until noted. The total RNA samples were treated according to the above SLAM-seq Kinetics Kit protocol. In brief, 35 μl of the IAA/OS/NP master mix, composed of 5 μl of freshly prepared Iodoacetamide (IAA, 100 mM), 25 μl of Organic Solvent (OS) and 5 μl of Sodium Phosphate (NP) was mixed with 15 μl of RNA (5 μg) for each sample. Reaction was incubated at 50°C for 15 mins before addition of 1 μl of Stopping Reagent (SR) and mixing to terminate the reaction. At this point, exposure to light was allowed. RNA was precipitated by adding 1 μl of Carrier Substance (CS), 5 μl of Sodium Acetate (NA) and 125 μl of 100% EtOH.

The purified RNAs (1 μg) from the above were mixed with Spike-ins (Qiagen, Cat# 331535) followed by small RNA library construction using Qiagen QIAseq miRNA Library Kit (Cat# 331502) according to the protocol. In brief, 3’ and 5’ adapters were ligated to the RNAs sequentially followed by reverse transcription using RT primers containing unique molecular index (UMI), cDNA cleanup, library amplification and library cleanup. This kit incorporates modified oligonucleotides to eliminate adapter dimers during the library construction and bead-based cleanups to eliminate unwanted background, bypassing the need for gel purification and size selection during library construction. The final libraries were sent for QC analysis and sequencing by Illumina PE100.

### Data analysis for small RNA SLAM-seq

Sequenced reads were analyzed with a Nextflow (Di Tommaso et al. 2017) pipeline that orchestrated public bioinformatics tools and custom python scripts based on rnalib v0.0.4 (Popitsch and Ameres 2024), pysam v0.23.0, pandas v2.2.3, numpy v1.26.4 and scipy v1.15.2 packages. Briefly, we first trimmed sequencing adapters from the reads (first mate only) with umitools v1.1.3 (Smith et al. 2017), matching for the QIAseq adapter sequence (AACTGTAGGCACCATCAAT) and extracting 12nt UMIs downstream. We then removed and counted reads that globally aligned to the 52 QIAseq spikein sequences using BioPython v1.85 (parameters: open_gap_score=-1 and defaults otherwise). Reads shorter than 15nt were dropped as too short. Reads that aligned with a length normalized alignment score of at least 0.9 were considered as spikein reads and assigned to the spikein sequence with the highest score. Remaining reads were then mapped to a transcriptome created from miRBase ‘miRNA_primary_transcript’ sequences (padded up- and downstream by 10nt; overlapping sequences were merged and combined to single contigs) using NextGenMap v0.5.5 (Sedlazeck et al. 2013) in SLAM-seq aware mode (parameters: --slam-seq 2 -very-sensitive -n 3 --strata --kmer 8 --kmer-skip 0 -s 0.0 --min-residues 18 --gap-read-penalty 50 --gap-ref-penalty 50 --gap-extend-penalty 50). We then removed soft-clipped sequences from the aligned reads (stemming, for example, from untemplated additions to the miRNAs) and then combined those with all unmapped reads from the previous step.

These pre-processed reads were then mapped to the human genome (GRCh38) using bowtie v 1.3.1 (parameters: -v 3 -k 100 --best --strata) (Langmead et al. 2009) and all reads with at least 1 mismatch that did not overlap with miRBase annotated miRNA loci were filtered as ‘off-target’ reads. Finally, we continued processing using the original trimmed sequences (i.e., including tailed sequences) and mapped again to the transcriptome using NextGenMap (parameters as above). The motivation for this off-target filtering procedure was to remove sequences from non-miRNA loci to avoid false-positive misalignments while retaining true positive miRNA reads with large edit distance (e.g., due to SLAM-seq induced T-to-C conversions, or due to tailing) to mature miRNA sequences.

We then de-duplicated those alignments based on the extracted UMI sequences using umitools dedup (parameters: --method unique), removed all reads with more than 10 mismatches as potential mapping artifacts and counted miRNA sequences using a custom Python script. We assigned strand-matching reads to pre-miRNAs if they either matched the miRBase annotated ‘miRNA_primary_transcript’ annotations (with a tolerance of 10nt up- or downstream) or if the whole read was enveloped by these annotations and covered at least 50% of those. We counted strand-matching reads as miRNA reads if their 5’-end matched the annotated miRNA 5’-end with a tolerance of 5nt up-/downstream, if their 3’-end matched the annotated 3’-end with a tolerance up 5nt up- and 15nt downstream and if the read covered at least 50% of the annotated miRNA interval. Before this check, we clipped prefix and postfix (tailing) bases at the first mismatch positions beyond the miRNA annotations. Counts of multimapping reads were normalized by the number of optimal alignment positions (1/NH).

Our small RNA libraries were prepared with synthetic spike-in controls, but analysis indicated variable spike-in recovery across wild type replicates (**Supplementary Table 1**, spikein_counts tab). Moreover, if normalizing to the apparently lower amount of spike-in reads in wild type libraries, this would suggest globally decreased miRNA levels in mutants. However, Northern blotting tests confirmed this was not the case, and rather that there were selective up-and downregulation of miRNAs when normalizing by total RNA amounts (**Supplementary Figure 9**). Consequently, spike-in normalization was not applied in downstream analyses. Instead, read counts were further normalized to sequencing depth (reads per million miRNA reads; RPM). In the clipped reads, we also called strand-specific T-to-C conversions and respective convertible positions (filtering for per-base qualities<20 and randomly skipping calls with a probability of 0.0006 that we estimated from mismatches in the sequencing data to account for randomly spread sequencing errors).

For calculating fractions of T-to-C conversions (FTC) per miRNA, we calculated the ratio of converted over convertible bases within the first 18 annotated nucleotides of a miRNA. For calculating effective 4sU (4sU_eff_) concentrations per sample, we fitted a zero-inflated binomial model (ZIB ∼ [0 with probability ψ and B(n; p) with probability 1 – ψ], where B is the binomial distribution with parameters n corresponding to the number of convertible positions, p corresponding to the probability of T-to-C conversions (4sU_eff_) and 1-ψ corresponding to the fraction of new reads (i.e., reads from miRNAs created after addition of 4sU). We fitted this model to the distribution of conversions per miRNA read with different numbers of convertible positions within the first annotated 18nt and excluding the first position due to an observed underrepresentation of conversions in our alignments. We fitted 10 times to subsamples of 50k reads, calculated the mean of the predicted binomial probabilities and then calculated the median of these values over a range of convertible position parameters (n in [4; 6] based on the data distributions; data not shown). We then used the fitted p (i.e., 4sU_eff_) values for estimating the fraction of new reads per miRNA: we fitted a ZIB model to the respective T-to-C conversion distributions per miRNA using the same procedure as above (subsample size 10k) while fixing the binomial probability to the estimated 4sU_eff_ values and extracting 1-ψ (i.e., frac_new).

To calculate counts of new reads, we multiplied these fractions with RPM normalized read count values. For calculating differential expression of pre-miRNA and miRNA reads, we used the pyDESeq2 package v0.5.2 (Muzellec et al. 2023), miRNAs were considered significantly DE if their adjusted P-value was < 0.05 and their absolute log2 fold change was > 1. The final set of considered miRNAs was determined by filtering for a minimum (raw) read count of 50 in any of the considered samples. Additionally, we removed 69 miRNAs with T-to-C convertible alignment columns within the 1st 18nt of the miRNA annotation that had a coverage > 10 and more than 75% converted bases as potential mapping artifacts. Our final dataset contains read count data for 713 miRNAs and fractions of new reads for 441 miRNAs; all final counts are in **Supplementary Table 1**. For comparison with *in vitro* Microprocessor pri-miRNA cleavage (Figure 6E), we downloaded Supplemental Table S6-2 comprising efficiency scores of pri-miRNA cleavage by recombinant Microprocessor+SRSF3 (Kim et al. 2021) and integrated with our dataset (**Supplementary Table 2**). All data was analyzed and plotted in iPython notebooks, data QC was conducted using miRTrace v1.0.1(Kang et al. 2018), samtools v1.18 (Danecek et al. 2021) and fastqc v0.11.9.

The raw SLAM-seq data were deposited at the European Nucleotide Archive under accession PRJEB101478. In addition, the raw pipeline results and analysis notebook for all the computational studies were assembled as a Zenodo repository available at 10.5281/zenodo.17464307.

## Acknowledgements

We thank Shuo Gu for providing *DGCR8-KO* and *Drosha/DGCR8-dKO* HEK293T cells used in the main figures, Wei Xie and Dinshaw Patel for advice on mutating the ERH dimerization interface, and Sebastian Herzog for providing SAFB1 and SAFB2 cDNAs used for cloning constructs. We thank the Developmental Studies Hybridoma Bank for providing antibodies to the community at cost. SLA lab was supported by the European Union (ERC, RiboTrace, CoG-866166), the Austrian Science Fund (FWF, 10.55776/F80 and 10.55776/DOC177), the Vienna Science and Technology Fund (WWTF, LS23-053), and the Austrian Academy of Sciences. Work in the ECL lab was supported by the National Institutes of Health (R01-GM083300), Tow Center for Developmental Oncology, and MSK Core Grant P30-CA008748.

## Author Contributions

R.S. conceived and designed the study. R.S. performed all the experiments and analyzed the data. S.L. analyzed genomic data. N.P. analyzed the small RNA SLAM-seq data under supervision of S.A. R.S. and E.C.L. wrote the manuscript, with input from all authors.

## Supplementary Figures

**Supplementary Figure 1. Summary of proposed models for miRNA cluster assistance.** At its heart, the ability of an optimal miRNA neighbor to enhance the nuclear biogenesis of an adjacent suboptimal miRNA hairpin implies that increased local concentration of Microprocessor can facilitate cleavage of pri-miRNA hairpin that lacks features of an optimal miRNA hairpin. This model is agnostic as to the contribution of other factors. Trans-acting factors ERH and SAFB1/2 can promote miRNA cluster assistance. Various protein-protein interactions amongst themselves and/or with Microprocessor components have been reported, which influence potential models as to their roles during miRNA biogenesis. In one set of models, ERH and SAFB1/2 might work separately, or together, during the transfer or stabilization of Microprocessor on the suboptimal miRNA neighbor. In another set of models ERH and SAFB1/2 acting separately, or in conjunction, might facilitate the formation of higher order Microprocessor complexes. In principle, these could form on the initial optimal miRNA hairpin, or bridge a pair of miRNA hairpins.

**Supplementary Figure 2. Genotyping of ERH, SAFB1/2 and *mir-3618* mutations.** Shown are the sequenced alleles of *ERH*, *SAFB1*, *SAFB2* indels and *mir-3618* deletions recovered in HEK293T cells used in this study.

**Supplementary Figure 3. Processing of *mir-144/451* or solo *mir-451* in cells depleted of Microprocessor cofactors.** (A) Comparison of miRNA processing by Northern blotting from HEK293T cells depleted of either of ERH, SAFB1, SAFB2 or their combinations. Co-transfected *mir-375* and endogenous let-7a and U6 snRNAs were probed as controls. RNA size markers (nt) are shown on the left. Microprocessing of suboptimal *pri-mir-451*, but not optimal *pri-mir-144* and *pri-mir-375*, is greatly repressed in *ERH* or *SAFB2* single mutant cells, while no defect was observed in *SAFB1* single mutant cells. But interestingly, *SAFB1/2* double mutant showed stronger repression on *pri-mir-451* processing than *SAFB2* single mutant, indicating a collaborative activity of SAFB1 and SAFB2. In contrast, absence of the neighboring *mir-144* hairpin almost completely repressed *mir-451* expression in all these cell lines. (B) Cross-rescue of *pri-mir-451* processing in mutant cells. While ERH cDNA can restore *mir-451* biogenesis, overexpressing SAFB2 also partially restored *mir-451* in *ERH* mutant cells. In contrast, only SAFB2, but not SAFB1 and ERH, can rescue *mir-451* biogenesis in *SAFB1/2* double mutant cells. These results further confirmed the unique and critical role of SAFB2 in cluster assistance.

**Supplementary Figure 4. Physical and function interactions between ERH and miRNA pathway factors.** (A) Co-immunoprecipitation (co-IP) assays of wildtype and variant ERH proteins bearing the designated mutations confirm that these mutants ablate dimerization of ERH. (B) Western blotting of DGCR8 variants lacking the N-terminal ERH interaction domain (residues 1-145) and/or that carry mutations in the DGCR8 dimerization domain confirm that all of these accumulate to similar levels in cells. (C) Northern blotting of rescue assays of DGCR8 variants transfected into DGCR8-KO cells. Deletion of the N-terminal domain does not affect rescue capacity of DGCR8 for optimal or suboptimal miRNAs, but loss of dimerization compromises its activity (especially for point mutant C352A). This experiment is similar to main Figure 3H, but uses an independent DGCR8-KO line generated in our lab.

**Supplementary Figure 5. Interaction tests between SAFB1/2 and Microprocessor factors.** HEK293T cells were transfected with the indicated tagged plasmids. HA-tagged factors were immunoprecipitated and Flag-tagged factors were tested for co-immunoprecipitation (co-IP). (A) The minimal fragment SAFB2[512-726] fully supports miRNA cluster assistance in SAFB1/2 mutant cells. Co-IP assays showed only mild, or no stable interactions between SAFB2[512-726] and full length Drosha, or Drosha deleted for its N-terminus (reported to contain the SAFB2 binding region). Assays were conducted in wildtype and knockout cells for both Microprocessor components. (B) Co-IP assays did not detect stable dimerization of functional SAFB2[512-726], and did not detect stable association of SAFB2[512-726] with ERH, either in wildtype cells or Microprocessor mutant cells. (C) Co-IP assays indicate that neither dimerization of DGCR8, nor stable formation of Microprocessor complex, requires SAFB1/2.

**Supplementary Figure 6. Processing and QC statistics of SLAM-seq data from Microprocessor cofactor mutants.** (A) Raw read counts before (all) and after demultiplexing (pass: reads for further processing, spikein: reads assigned to spike-in sequences, read_too_short: filtered reads). (B) Median read lengths of the respective groups (error bar indices 95% ci over all datasets). The grey band highlights the expected size of mature miRNAs (20-24 nts). (C) Deduplication statistics. mapped_reads: number of alignments before deduplication. dedup_reads: number of reads after deduplication. (D) Read fate statistics from our small RNA analysis pipeline, error bars indicate variation between samples. Final miRNA counting was based on deduplicated alignments. (E) Size distribution of SLAM-seq libraries from 6hr labeling of wt, ERH, SAFB1/2 and ERH+SAFB1/2 HEK293T cells. Filtering was done to remove potentially off-target mappings. Of the pass reads (“final” columns), most reads are in the size range of mature miRNAs, but there are also minor populations of pre-miRNA-sized reads. (F) RNA types of reads from the pass and off-target populations. (G) miRNA complexity of the pass and off-target populations. (H) The replicate libraries are well-correlated.

**Supplementary Figure 7. Analysis of miRNAs previously assessed for dependency on Microprocessor cofactors.** (A) Shown are the miRNAs reported by Kwon et al Nucleic Acids Research 2020 (Figure 7) to be dependent on ERH and/or SAFB1/2 factors, using different cell models. For ease of comparison, we plotted these in the same order as in the prior study, using our data from ERH, SAFB1/2 and ERH+SAFB1/2 mutant HEK293T cells, and recorded their clustered status with circles or dots (to the right of the heatmap). For the most part, our data agree with the prior study, except that let-7e-5p was mildly downregulated in our data (as opposed to upregulated in ERH-kd), and a few other miRNAs were mildly downregulated in our data (but upregulated in prior ERH-kd data).

**Supplementary Figure 8. Volcano plots of miRNA dysregulation in mutants of Microprocessor cofactors.** Volcano plots of differential miRNA expression in the indicated mutant HEK293T backgrounds, compared to wildtype cells. The plot contains all miRNAs with sufficient expression (n=530), and a negative log2 fold change indicates downregulation in the respective mutant condition. Icons and colors indicate clustered status (X) and validated suboptimal miRNAs (square).

**Supplementary Figure 9. Validation of endogenous miRNAs in different genotypes by Northern blots.** Total RNAs of WT, ERH and/or SAFB1/2 mutant HEK293T cells (the same batch of RNAs used for SLAM-seq) were run on three blots to probe different miRNAs that (A) are preferentially downregulated in SAFB1/2 compared to ERH, (B) are dependent on both ERH and SAFB1/2, or that (C) are upregulated in ERH and SAFB1/2 mutants. miR-16 is an optimal miRNA that is relatively unchanged in mutants and deep sequencing, and was used as control.

**Supplementary Figure 10. UpSet plots of miRNA dysregulation in mutants of Microprocessor cofactors.** Shown are UpSet plots of differentially expressed miRNAs including all (left panels) and only clustered miRNA loci (right panels). These were divided according to plots of all miRNAs (top panels), only upregulated miRNAs (middle panels) and only downregulated miRNAs (bottom panels). It can be seen that SAFB1/2 and ERH+SAFB1/2 mutants account for the majority of downregulated miRNAs, and that a substantial portion of these are also shared by ERH mutants. In addition, amongst clustered miRNAs, there are far more that are downregulated than upregulated in these mutants, consistent with the notion that disruption of cluster assistance is a major drive of miRNA dysregulation in these mutants, as opposed to other direct roles or indirect consequences of these mutant cells.

**Supplementary Figure 11. Additional examples of suboptimal miRNAs with dependency on ERH and/or SAFB1/2.** We extracted illustrative miRNAs with preferential downregulation in ERH mutant, in SAFB1/2 mutant (without or with additional ERH depletion), or in combined ERH+SAFB1/2 mutant HEK293T cells. Note that as SAFB1/2 depletion had the largest effect on miRNA dysregulation in these datasets, we were able to select examples that show more substantial defects in miRNA expression than in other cases. For example, there are few miRNAs that show an enhanced downregulation in the combined triple mutant data. For all genotypes, we highlighted example loci where similar trends were observed in both mature miRNA and passenger strand (miRNA*) species of the given miRNA hairpin, which helped support the notion that these reflect biogenesis defects (as opposed to stability effects). We show predicted secondary structures for these loci. In many cases, they exhibit predicted suboptimal stems that may underlie their sensitivity to Microprocessor cofactors. Amongst ERH-sensitive miRNAs, many exhibit seemingly typical pri-miRNA structures. It remains to be seen if these are indeed hairpin suboptimality that is not reflected by structure predictions, or if these ERH-dependent miRNAs involve effects other than cluster assistance.

**Supplementary Figure 12. Quality control assessment of newly-synthesized miRNAs in SLAM-seq data.** SLAM-seq libraries were constructed following labeling with 4sU for 6 hours. Fractions of reads from newly synthesized miRNAs were estimated from the observed T-to-C conversion patterns as described in Methods. (A) Amongst total reads, mature miRNA strands dominate all the libraries for the four genotypes. (B) However, amongst newly-synthesized reads, miRNA* (passenger strands) are heavily enriched. This behavior is expected, since miRNA* species are degraded following their ejection from the mature Argonaute-miRNA complex. This assessment supports that the SLAM-seq protocol indeed captured newly-synthesized small RNAs (transcribed within the 6 hr labeling period).

**Supplementary Figure 13. Additional Northern blotting of *DGCR8* miRNAs.** (A) Northern blotting shows that endogenous *pre-mir-3618* is readily detected, and is lost in *mir-3618* deletion HEK293T cells. However, no mature miR-3618 is visible. Other probings show mature miR-92a-3p and let-7a-5p. Note that these are the same original blots that are cropped in main Figure 7C to highlight the specific bands. (B) Ectopic expression of DGCR8 miRNA constructs containing *mir-3618*, *mir-1306*, or both, increase signals for *pre-mir-3618*, but still do not reveal mature miR-3618.

## References

Aguado LC, Schmid S, May J, Sabin LR, Panis M, Blanco-Melo D, Shim JV, Sachs D, Cherry S, Simon AE et al. 2017. RNase III nucleases from diverse kingdoms serve as antiviral effectors. Nature 547: 114–117. 10.1038/nature22990

Arai R, Kukimoto-Niino M, Uda-Tochio H, Morita S, Uchikubo-Kamo T, Akasaka R, Etou Y, Hayashizaki Y, Kigawa T, Terada T et al. 2005. Crystal structure of an enhancer of rudimentary homolog (ERH) at 2.1 Angstroms resolution. Protein Sci 14: 1888–1893. 10.1110/ps.051484505

Aschenwald S, Panda A, Wurzer T, Baumgartner S, Ertl V, Hatzer J, Villunger A, Falk S, Herzog S. 2025. Dual function of ERH in primary miRNA biogenesis. BioRxiv: https://www.biorxiv.org/content/10.1101/2025.1109.1123.678008v678001.

Auyeung VC, Ulitsky I, McGeary SE, Bartel DP. 2013. Beyond secondary structure: primary-sequence determinants license pri-miRNA hairpins for processing. Cell 152: 844–858. 10.1016/j.cell.2013.01.031

Banko MI, Krzyzanowski MK, Turcza P, Maniecka Z, Kulis M, Kozlowski P. 2013. Identification of amino acid residues of ERH required for its recruitment to nuclear speckles and replication foci in HeLa cells. PloS one 8: e74885. 10.1371/journal.pone.0074885

Barr I, Smith AT, Chen Y, Senturia R, Burstyn JN, Guo F. 2012. Ferric, not ferrous, heme activates RNA-binding protein DGCR8 for primary microRNA processing. Proceedings of the National Academy of Sciences of the United States of America 109: 1919–1924. 10.1073/pnas.1114514109

Barr I, Smith AT, Senturia R, Chen Y, Scheidemantle BD, Burstyn JN, Guo F. 2011. DiGeorge critical region 8 (DGCR8) is a double-cysteine-ligated heme protein. The Journal of biological chemistry 286: 16716–16725. 10.1074/jbc.M110.180844

Bartel DP. 2018. Metazoan MicroRNAs. Cell 173: 20–51. 10.1016/j.cell.2018.03.006

Bartel DP, Chen CZ. 2004. Micromanagers of gene expression: the potentially widespread influence of metazoan microRNAs. Nature reviews Genetics 5: 396–400. 10.1038/nrg1328

Berezikov E, Chung WJ, Willis J, Cuppen E, Lai EC. 2007. Mammalian mirtron genes. Molecular cell 28: 328–336. 10.1016/j.molcel.2007.09.028

Bogerd HP, Whisnant AW, Kennedy EM, Flores O, Cullen BR. 2014. Derivation and characterization of Dicer- and microRNA-deficient human cells. RNA 20: 923–937. 10.1261/rna.044545.114

Brennecke J, Stark A, Russell RB, Cohen SM. 2005. Principles of microRNA-target recognition. PLoS biology 3: e85. 10.1371/journal.pbio.0030085

Cheloufi S, Dos Santos CO, Chong MM, Hannon GJ. 2010. A dicer-independent miRNA biogenesis pathway that requires Ago catalysis. Nature 465: 584–589. 10.1038/nature09092

Cherney RE, Eberhard QE, Giri G, Mills CA, Porrello A, Zhang Z, White D, Trotman JB, Herring LE, Dominguez D et al. 2023. SAFB associates with nascent RNAs and can promote gene expression in mouse embryonic stem cells. RNA 29: 1535–1556. 10.1261/rna.079569.122

Cifuentes D, Xue H, Taylor DW, Patnode H, Mishima Y, Cheloufi S, Ma E, Mane S, Hannon GJ, Lawson ND et al. 2010. A novel miRNA processing pathway independent of Dicer requires Argonaute2 catalytic activity. Science 328: 1694–1698. 10.1126/science.1190809

Clarke AW, Hoye E, Hembrom AA, Paynter VM, Vinther J, Wyrozemski L, Biryukova I, Formaggioni A, Ovchinnikov V, Herlyn H et al. 2025. MirGeneDB 3.0: improved taxonomic sampling, uniform nomenclature of novel conserved microRNA families and updated covariance models. Nucleic acids research 53: D116–D128. 10.1093/nar/gkae1094

Cordeiro Rodrigues RJ, de Jesus Domingues AM, Hellmann S, Dietz S, de Albuquerque BFM, Renz C, Ulrich HD, Sarkies P, Butter F, Ketting RF. 2019. PETISCO is a novel protein complex required for 21U RNA biogenesis and embryonic viability. Genes & development 33: 857–870. 10.1101/gad.322446.118

Dai L, Chen K, Youngren B, Kulina J, Yang A, Guo Z, Li J, Yu P, Gu S. 2016. Cytoplasmic Drosha activity generated by alternative splicing. Nucleic acids research 44: 10454–10466. 10.1093/nar/gkw668

Danecek P, Bonfield JK, Liddle J, Marshall J, Ohan V, Pollard MO, Whitwham A, Keane T, McCarthy SA, Davies RM et al. 2021. Twelve years of SAMtools and BCFtools. Gigascience 10. 10.1093/gigascience/giab008

Di Tommaso P, Chatzou M, Floden EW, Barja PP, Palumbo E, Notredame C. 2017. Nextflow enables reproducible computational workflows. Nature biotechnology 35: 316–319. 10.1038/nbt.3820

Drakouli S, Lyberopoulou A, Papathanassiou M, Mylonis I, Georgatsou E. 2017. Enhancer of rudimentary homologue interacts with scaffold attachment factor B at the nuclear matrix to regulate SR protein phosphorylation. FEBS J 284: 2482–2500. 10.1111/febs.14141

Faller M, Matsunaga M, Yin S, Loo JA, Guo F. 2007. Heme is involved in microRNA processing. Nature structural & molecular biology 14: 23–29. 10.1038/nsmb1182

Fang W, Bartel DP. 2015. The Menu of Features that Define Primary MicroRNAs and Enable De Novo Design of MicroRNA Genes. Molecular cell 60: 131–145. 10.1016/j.molcel.2015.08.015

Fang W, Bartel DP. 2020. MicroRNA Clustering Assists Processing of Suboptimal MicroRNA Hairpins through the Action of the ERH Protein. Molecular cell 78: 289–302 e286. 10.1016/j.molcel.2020.01.026

Forcella P, Ifflander N, Rolando C, Balta EA, Lampada A, Giachino C, Mukhtar T, Bock T, Taylor V. 2024. SAFB regulates hippocampal stem cell fate by targeting Drosha to destabilize Nfib mRNA. eLife 13. 10.7554/eLife.74940

Garg A, Shang R, Cvetanovic T, Lai EC, Joshua-Tor L. 2024. The structural landscape of Microprocessor-mediated processing of pri-let-7 miRNAs. Molecular cell 84: 4175–4190 e4176. 10.1016/j.molcel.2024.09.008

Haar J, Contrant M, Bernhardt K, Feederle R, Diederichs S, Pfeffer S, Delecluse HJ. 2016. The expression of a viral microRNA is regulated by clustering to allow optimal B cell transformation. Nucleic acids research 44: 1326–1341. 10.1093/nar/gkv1330

Han J, Lee Y, Yeom KH, Nam JW, Heo I, Rhee JK, Sohn SY, Cho Y, Zhang BT, Kim VN. 2006. Molecular basis for the recognition of primary microRNAs by the Drosha-DGCR8 complex. Cell 125: 887–901. 10.1016/j.cell.2006.03.043

Han J, Pedersen JS, Kwon SC, Belair CD, Kim YK, Yeom KH, Yang WY, Haussler D, Blelloch R, Kim VN. 2009. Posttranscriptional crossregulation between Drosha and DGCR8. Cell 136: 75–84. 10.1016/j.cell.2008.10.053

Hazra D, Andric V, Palancade B, Rougemaille M, Graille M. 2020. Formation of S. pombe Erh1 homodimer mediates gametogenic gene silencing and meiosis progression. Scientific reports 10: 1034. 10.1038/s41598-020-57872-4

Herzog VA, Reichholf B, Neumann T, Rescheneder P, Bhat P, Burkard TR, Wlotzka W, von Haeseler A, Zuber J, Ameres SL. 2017. Thiol-linked alkylation of RNA to assess expression dynamics. Nature methods 14: 1198–1204. 10.1038/nmeth.4435

Hong Y, Bie L, Zhang T, Yan X, Jin G, Chen Z, Wang Y, Li X, Pei G, Zhang Y et al. 2024. SAFB restricts contact domain boundaries associated with L1 chimeric transcription. Molecular cell 84: 1637–1650 e1610. 10.1016/j.molcel.2024.03.021

Hutter K, Lohmuller M, Jukic A, Eichin F, Avci S, Labi V, Szabo TG, Hoser SM, Huttenhofer A, Villunger A et al. 2020. SAFB2 Enables the Processing of Suboptimal Stem-Loop Structures in Clustered Primary miRNA Transcripts. Molecular cell 78: 876–889 e876. 10.1016/j.molcel.2020.05.011

Ilik IA, Glazar P, Tse K, Brandl B, Meierhofer D, Muller FJ, Smith ZD, Aktas T. 2024. Autonomous transposons tune their sequences to ensure somatic suppression. Nature 626: 1116–1124. 10.1038/s41586-024-07081-0

Jang H, Park J, Kim VN. 2025. ERH promotes primary microRNA processing beyond cluster assistance. Nature communications 16: 7913. 10.1038/s41467-025-63015-y

Jin W, Wang J, Liu CP, Wang HW, Xu RM. 2020. Structural Basis for pri-miRNA Recognition by Drosha. Molecular cell 78: 423–433 e425. 10.1016/j.molcel.2020.02.024

Kadener S, Rodriguez J, Abruzzi KC, Khodor YL, Sugino K, Marr MT, 2nd, Nelson S, Rosbash M. 2009. Genome-wide identification of targets of the drosha-pasha/DGCR8 complex. RNA 15: 537–545. 10.1261/rna.1319309

Kang W, Eldfjell Y, Fromm B, Estivill X, Biryukova I, Friedlander MR. 2018. miRTrace reveals the organismal origins of microRNA sequencing data. Genome biology 19: 213. 10.1186/s13059-018-1588-9

Kavanaugh G, Zhao R, Guo Y, Mohni KN, Glick G, Lacy ME, Hutson MS, Ascano M, Cortez D. 2015. Enhancer of Rudimentary Homolog Affects the Replication Stress Response through Regulation of RNA Processing. Molecular and cellular biology 35: 2979–2990. 10.1128/MCB.01276-14

Kim H, Lee YY, Kim VN. 2025. The biogenesis and regulation of animal microRNAs. Nature reviews Molecular cell biology 26: 276–296. 10.1038/s41580-024-00805-0

Kim K, Baek SC, Lee YY, Bastiaanssen C, Kim J, Kim H, Kim VN. 2021. A quantitative map of human primary microRNA processing sites. Molecular cell 81: 3422–3439 e3411. 10.1016/j.molcel.2021.07.002

Korn SM, Von Ehr J, Dhamotharan K, Tants JN, Abele R, Schlundt A. 2023. Insight into the Structural Basis for Dual Nucleic Acid-Recognition by the Scaffold Attachment Factor B2 Protein. Int J Mol Sci 24. 10.3390/ijms24043286

Kozlowski P. 2023. Thirty Years with ERH: An mRNA Splicing and Mitosis Factor Only or Rather a Novel Genome Integrity Protector? Cells 12. 10.3390/cells12202449

Kretov DA, Walawalkar IA, Mora-Martin A, Shafik AM, Moxon S, Cifuentes D. 2020. Ago2-Dependent Processing Allows miR-451 to Evade the Global MicroRNA Turnover Elicited during Erythropoiesis. Molecular cell 78: 317–328 e316. 10.1016/j.molcel.2020.02.020

Kwon SC, Baek SC, Choi YG, Yang J, Lee YS, Woo JS, Kim VN. 2019. Molecular Basis for the Single-Nucleotide Precision of Primary microRNA Processing. Molecular cell 73: 505–518 e505. 10.1016/j.molcel.2018.11.005

Kwon SC, Jang H, Shen S, Baek SC, Kim K, Yang J, Kim J, Kim JS, Wang S, Shi Y et al. 2020. ERH facilitates microRNA maturation through the interaction with the N-terminus of DGCR8. Nucleic acids research 48: 11097–11112. 10.1093/nar/gkaa827

Kwon SC, Nguyen TA, Choi YG, Jo MH, Hohng S, Kim VN, Woo JS. 2016. Structure of Human DROSHA. Cell 164: 81–90. 10.1016/j.cell.2015.12.019

Lai EC. 2002. Micro RNAs are complementary to 3’ UTR sequence motifs that mediate negative post-transcriptional regulation. Nature genetics 30: 363–364. 10.1038/ng865

Lai EC, Burks C, Posakony JW. 1998. The K box, a conserved 3’ UTR sequence motif, negatively regulates accumulation of enhancer of split complex transcripts. Development 125: 4077–4088. 10.1242/dev.125.20.4077

Lai EC, Posakony JW. 1997. The Bearded box, a novel 3’ UTR sequence motif, mediates negative post-transcriptional regulation of *Bearded* and *Enhancer of split* Complex gene expression. Development 124: 4847–4856. 10.1242/dev.124.23.4847

Langmead B, Trapnell C, Pop M, Salzberg SL. 2009. Ultrafast and memory-efficient alignment of short DNA sequences to the human genome. Genome biology 10: R25. 10.1186/gb-2009-10-3-r25

Lee D, Nam JW, Shin C. 2017. DROSHA targets its own transcript to modulate alternative splicing. RNA 23: 1035–1047. 10.1261/rna.059808.116

Lee RC, Feinbaum RL, Ambros V. 1993. The *C. elegans* heterochronic gene *lin-4* encodes small RNAs with antisense complementarity to *lin-14*. Cell 75: 843–854. 10.1016/0092-8674(93)90529-y

Lee YY, Kim H, Kim VN. 2023. Sequence determinant of small RNA production by DICER. Nature 615: 323–330. 10.1038/s41586-023-05722-4

Lewis BP, Shih IH, Jones-Rhoades MW, Bartel DP, Burge CB. 2003. Prediction of mammalian microRNA targets. Cell 115: 787–798. 10.1016/s0092-8674(03)01018-3

Ma H, Wu Y, Choi JG, Wu H. 2013. Lower and upper stem-single-stranded RNA junctions together determine the Drosha cleavage site. Proceedings of the National Academy of Sciences of the United States of America 110: 20687–20692. 10.1073/pnas.1311639110

Maurin T, Cazalla D, Yang S, Jr., Bortolamiol-Becet D, Lai EC. 2012. RNase III-independent microRNA biogenesis in mammalian cells. RNA 18: 2166–2173. 10.1261/rna.036194.112

McGeary SE, Lin KS, Shi CY, Pham TM, Bisaria N, Kelley GM, Bartel DP. 2019. The biochemical basis of microRNA targeting efficacy. Science 366. 10.1126/science.aav1741

Mechtler P, Johnson S, Slabodkin H, Cohanim AB, Brodsky L, Kandel ES. 2017. The evidence for a microRNA product of human DROSHA gene. RNA biology 14: 1508–1513. 10.1080/15476286.2017.1342934

Mohammed J, Siepel A, Lai EC. 2014. Diverse modes of evolutionary emergence and flux of conserved microRNA clusters. RNA 20: 1850–1863. 10.1261/rna.046805.114

Muzellec B, Telenczuk M, Cabeli V, Andreux M. 2023. PyDESeq2: a python package for bulk RNA-seq differential expression analysis. Bioinformatics 39. 10.1093/bioinformatics/btad547

Nguyen TA, Jo MH, Choi YG, Park J, Kwon SC, Hohng S, Kim VN, Woo JS. 2015. Functional Anatomy of the Human Microprocessor. Cell 161: 1374–1387. 10.1016/j.cell.2015.05.010

Norman M, Rivers C, Lee YB, Idris J, Uney J. 2016. The increasing diversity of functions attributed to the SAFB family of RNA-/DNA-binding proteins. The Biochemical journal 473: 4271–4288. 10.1042/BCJ20160649

Nussbacher JK, Yeo GW. 2018. Systematic Discovery of RNA Binding Proteins that Regulate MicroRNA Levels. Molecular cell 69: 1005–1016 e1007. 10.1016/j.molcel.2018.02.012

Okamura K, Hagen JW, Duan H, Tyler DM, Lai EC. 2007. The mirtron pathway generates microRNA-class regulatory RNAs in Drosophila. Cell 130: 89–100. 10.1016/j.cell.2007.06.028

Partin AC, Zhang K, Jeong BC, Herrell E, Li S, Chiu W, Nam Y. 2020. Cryo-EM Structures of Human Drosha and DGCR8 in Complex with Primary MicroRNA. Molecular cell 78: 411–422 e414. 10.1016/j.molcel.2020.02.016

Perez-Borrajero C, Podvalnaya N, Holleis K, Lichtenberger R, Karaulanov E, Simon B, Basquin J, Hennig J, Ketting RF, Falk S. 2021. Structural basis of PETISCO complex assembly during piRNA biogenesis in C. elegans. Genes & development 35: 1304–1323. 10.1101/gad.348648.121

Pogge von Strandmann E, Senkel S, Ryffel GU. 2001. ERH (enhancer of rudimentary homologue), a conserved factor identical between frog and human, is a transcriptional repressor. Biological chemistry 382: 1379–1385. 10.1515/BC.2001.170

Popitsch N, Ameres SL. 2024. Rnalib: a Python library for custom transcriptomics analyses. Bioinformatics 41. 10.1093/bioinformatics/btae751

Reichholf B, Herzog VA, Fasching N, Manzenreither RA, Sowemimo I, Ameres SL. 2019. Time-Resolved Small RNA Sequencing Unravels the Molecular Principles of MicroRNA Homeostasis. Molecular cell 75: 756–768 e757. 10.1016/j.molcel.2019.06.018

Renz A, Fackelmayer FO. 1996. Purification and molecular cloning of the scaffold attachment factor B (SAF-B), a novel human nuclear protein that specifically binds to S/MAR-DNA. Nucleic acids research 24: 843–849. 10.1093/nar/24.5.843

Rivers C, Idris J, Scott H, Rogers M, Lee YB, Gaunt J, Phylactou L, Curk T, Campbell C, Ule J et al. 2015. iCLIP identifies novel roles for SAFB1 in regulating RNA processing and neuronal function. BMC biology 13: 111. 10.1186/s12915-015-0220-7

Ruby JG, Jan CH, Bartel DP. 2007. Intronic microRNA precursors that bypass Drosha processing. Nature 448: 83–86. 10.1038/nature05983

Sedlazeck FJ, Rescheneder P, von Haeseler A. 2013. NextGenMap: fast and accurate read mapping in highly polymorphic genomes. Bioinformatics 29: 2790–2791. 10.1093/bioinformatics/btt468

Sergeant KA, Bourgeois CF, Dalgliesh C, Venables JP, Stevenin J, Elliott DJ. 2007. Alternative RNA splicing complexes containing the scaffold attachment factor SAFB2. Journal of cell science 120: 309–319. 10.1242/jcs.03344

Shang R, Baek SC, Kim K, Kim B, Kim VN, Lai EC. 2020. Genomic Clustering Facilitates Nuclear Processing of Suboptimal Pri-miRNA Loci. Molecular cell 78: 303–316 e304. 10.1016/j.molcel.2020.02.009

Shang R, Kretov DA, Adamson SI, Treiber T, Treiber N, Vedanayagam J, Chuang JH, Meister G, Cifuentes D, Lai EC. 2022. Regulated dicing of pre-mir-144 via reshaping of its terminal loop. Nucleic acids research 50: 7637–7654. 10.1093/nar/gkac568

Shang R, Lai EC. 2023. Parameters of clustered suboptimal miRNA biogenesis. Proceedings of the National Academy of Sciences of the United States of America 120: e2306727120. 10.1073/pnas.2306727120

Shang R, Lee S, Senavirathne G, Lai EC. 2023. microRNAs in action: biogenesis, function and regulation. Nature reviews Genetics 24: 816–833. 10.1038/s41576-023-00611-y

Shang R, Zhang F, Xu B, Xi H, Zhang X, Wang W, Wu L. 2015. Ribozyme-enhanced single-stranded Ago2-processed interfering RNA triggers efficient gene silencing with fewer off-target effects. Nature communications 6: 8430. 10.1038/ncomms9430

Smibert P, Bejarano F, Wang D, Garaulet DL, Yang JS, Martin R, Bortolamiol-Becet D, Robine N, Hiesinger PR, Lai EC. 2011. A Drosophila genetic screen yields allelic series of core microRNA biogenesis factors and reveals post-developmental roles for microRNAs. RNA 17: 1997–2010. 10.1261/rna.2983511

Smith T, Heger A, Sudbery I. 2017. UMI-tools: modeling sequencing errors in Unique Molecular Identifiers to improve quantification accuracy. Genome research 27: 491–499. 10.1101/gr.209601.116

Street LA, Rothamel KL, Brannan KW, Jin W, Bokor BJ, Dong K, Rhine K, Madrigal A, Al-Azzam N, Kim JK et al. 2024. Large-scale map of RNA-binding protein interactomes across the mRNA life cycle. Molecular cell 84: 3790–3809 e3798. 10.1016/j.molcel.2024.08.030

Townson SM, Dobrzycka KM, Lee AV, Air M, Deng W, Kang K, Jiang S, Kioka N, Michaelis K, Oesterreich S. 2003. SAFB2, a new scaffold attachment factor homolog and estrogen receptor corepressor. The Journal of biological chemistry 278: 20059–20068. 10.1074/jbc.M212988200

Treiber T, Treiber N, Plessmann U, Harlander S, Daiss JL, Eichner N, Lehmann G, Schall K, Urlaub H, Meister G. 2017. A Compendium of RNA-Binding Proteins that Regulate MicroRNA Biogenesis. Molecular cell 66: 270–284 e213. 10.1016/j.molcel.2017.03.014

Triboulet R, Chang HM, Lapierre RJ, Gregory RI. 2009. Post-transcriptional control of DGCR8 expression by the Microprocessor. RNA 15: 1005–1011. 10.1261/rna.1591709

Truscott M, Islam AB, Frolov MV. 2016. Novel regulation and functional interaction of polycistronic miRNAs. RNA 22: 129–138. 10.1261/rna.053264.115

Vilimova M, Contrant M, Randrianjafy R, Dumas P, Elbasani E, Ojala PM, Pfeffer S, Fender A. 2021. Cis regulation within a cluster of viral microRNAs. Nucleic acids research 49: 10018–10033. 10.1093/nar/gkab731

Wan C, Tempel W, Liu ZJ, Wang BC, Rose RB. 2005. Structure of the conserved transcriptional repressor enhancer of rudimentary homolog. Biochemistry 44: 5017–5023. 10.1021/bi047785w

Westholm JO, Lai EC. 2011. Mirtrons: microRNA biogenesis via splicing. Biochimie 93: 1897–1904. 10.1016/j.biochi.2011.06.017

Wightman B, Ha I, Ruvkun G. 1993. Posttranscriptional regulation of the heterochronic gene *lin-14* by *lin-4* mediates temporal pattern formation in *C. elegans*. Cell 75: 855–862. 10.1016/0092-8674(93)90530-4

Wojcik E, Murphy AM, Fares H, Dang-Vu K, Tsubota SI. 1994. Enhancer of rudimentaryp1, e(r)p1, a highly conserved enhancer of the rudimentary gene. Genetics 138: 1163–1170. 10.1093/genetics/138.4.1163

Xie M, Li M, Vilborg A, Lee N, Shu MD, Yartseva V, Sestan N, Steitz JA. 2013. Mammalian 5’-capped microRNA precursors that generate a single microRNA. Cell 155: 1568–1580. 10.1016/j.cell.2013.11.027

Yang JS, Lai EC. 2011. Alternative miRNA biogenesis pathways and the interpretation of core miRNA pathway mutants. Molecular cell 43: 892–903. 10.1016/j.molcel.2011.07.024

Yang JS, Maurin T, Robine N, Rasmussen KD, Jeffrey KL, Chandwani R, Papapetrou EP, Sadelain M, O’Carroll D, Lai EC. 2010. Conserved vertebrate mir-451 provides a platform for Dicer-independent, Ago2-mediated microRNA biogenesis. Proceedings of the National Academy of Sciences of the United States of America 107: 15163–15168. 10.1073/pnas.1006432107

Yoda M, Cifuentes D, Izumi N, Sakaguchi Y, Suzuki T, Giraldez AJ, Tomari Y. 2013. Poly(A)-specific ribonuclease mediates 3’-end trimming of Argonaute2-cleaved precursor microRNAs. Cell reports 5: 715–726. 10.1016/j.celrep.2013.09.029

Zamudio JR, Kelly TJ, Sharp PA. 2014. Argonaute-bound small RNAs from promoter-proximal RNA polymerase II. Cell 156: 920–934. 10.1016/j.cell.2014.01.041

Zeng Y, Cullen BR. 2005. Efficient processing of primary microRNA hairpins by Drosha requires flanking nonstructured RNA sequences. The Journal of biological chemistry 280: 27595–27603. 10.1074/jbc.M504714200

Zeng Y, Yi R, Cullen BR. 2005. Recognition and cleavage of primary microRNA precursors by the nuclear processing enzyme Drosha. The EMBO journal 24: 138–148. 10.1038/sj.emboj.7600491

Zhang X, Zeng Y. 2010. The terminal loop region controls microRNA processing by Drosha and Dicer. Nucleic acids research 38: 7689–7697. 10.1093/nar/gkq645

