## Supplementary Figures 1-13 for "Separable roles for Microprocessor and its cofactors ERH and SAFB1/2 during microRNA cluster assistance"

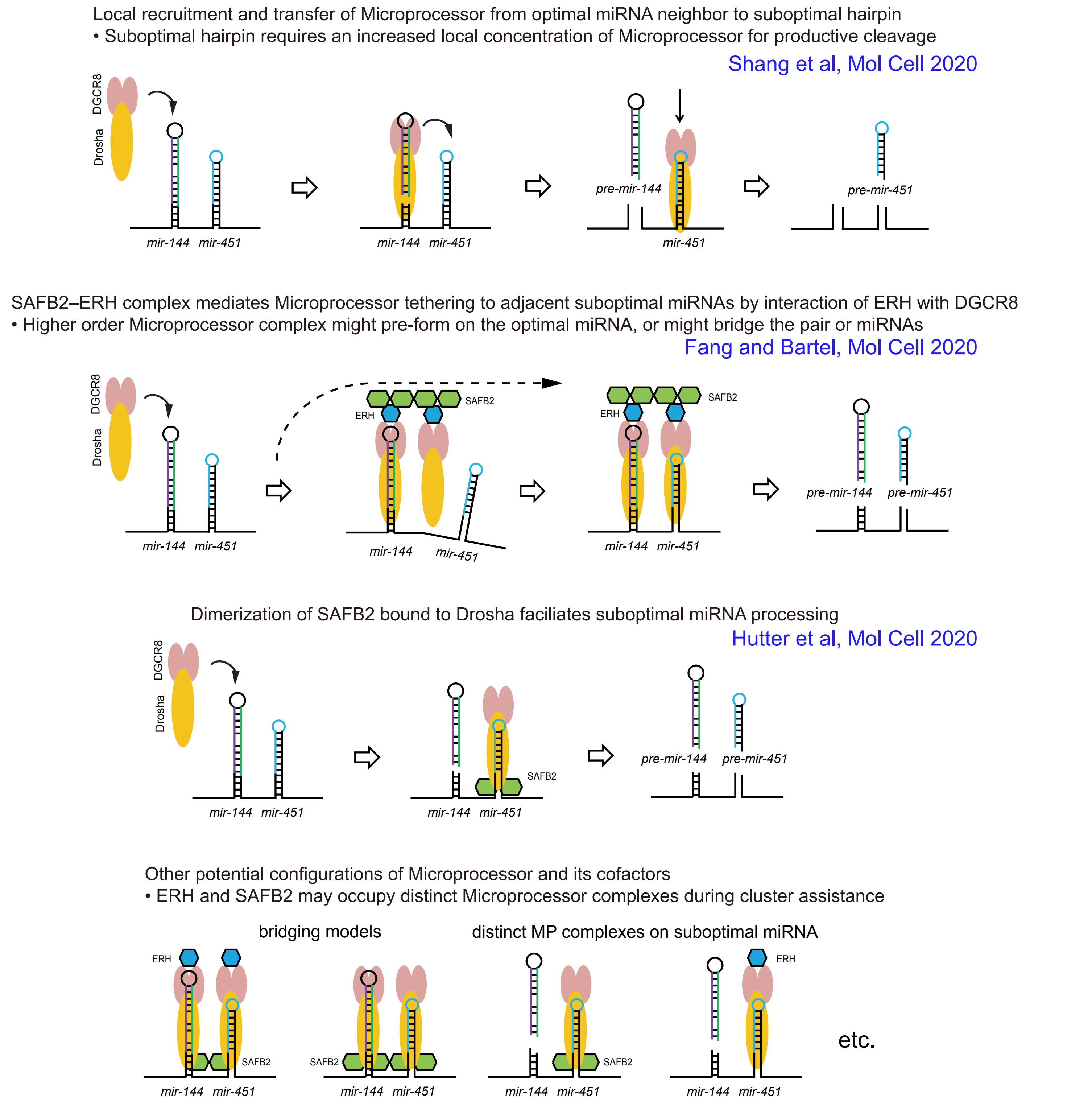


Supplementary Figure 1. Summary of proposed and plausible models for miRNA cluster assistance.
At its heart, the ability of an optimal miRNA neighbor to enhance the nuclear biogenesis of an adjacent suboptimal miRNA hairpin implies that increased local concentration of Microprocessor can facilitate cleavage of pri-miRNA hairpin that lacks features of an optimal miRNA hairpin. This model is agnostic as to the contribution of other factors. Trans-acting factors ERH and SAFB1/2 can promote miRNA cluster assistance. Various protein-protein interactions amongst themselves and/or with Microprocessor components have been reported, which influence potential models as to their roles during miRNA biogenesis. In one set of models, ERH and SAFB1/2 might work separately, or together, during the transfer or stabilization of Microprocessor on the suboptimal miRNA neighbor. In another set of models ERH and SAFB1/2 acting separately, or in conjunction, might facilitate the formation of higher order Microprocessor complexes. In principle, these could form on the initial optimal miRNA hairpin, or bridge a pair of miRNA hairpins.

Shang et al,
Supplementary Figure 1

Yellow highlighted sequences are NGG motifs for Cas9 recognition.

```
ERH(+/-) mutant#2
wt:  TCTCACACCATTTTGCTGGTACAGCCTACC AAGAGGCCAGAAGGCAGAACTTATGCTGACTACGAATCTGTGAATGAATGCATGGAAG
mut: TCTCACACCATTTTGCTGGTACAGCCTACCCAAGAGGCCAGAAGGCAGAACTTATGCTGACTACGAATCTGTGAATGAATGCATGGAAG  +1

SAFB1(+/-) mutant#4
wt:  GCAATTGAAGATGAAGGTGGTAATCCTGACGAAATTGAAATTACCTCCGAGGGAAACAAGAAAACATCAAA GAGGTCTAGCAAAG
mut: GCAATTGAAGATGAAGGTGGTAATCCTGACGAAATTGAAATTACCTCCGAGGGAAACAAGAAAACATCAAAAGAGGTCTAGCAAAG  +1

SAFB2(-/-) mutant#7
mut1: GCGGTTAAAGAAGAGGGGCAA             TTGGCATCGAGTTAGAAGCCACCAGCAAGAAGTCAGCCAAGAGATGTGTTAAAG  -13
mut2: GCGGTTAAAGAAGAGGGGCAAGATCCTGATG AATTGGCATCGAGTTAGAAGCCACCAGCAAGAAGTCAGCCAAGAGATGTGTTAAAG  -1

SAFB1(+/-)/SAFB2(-/-) mutant#14
SAFB1mut1: GCTAGAGCCGGCAGTTGAGCAGAGTAGTGCGGCCTCCGAGCTCGCGGAGGCCTCTAAGCGAGGAGCTCGCAGAAGCACCCACG  +1
SAFB1mut2: GCTAGAGCCGGCAGTTGAGCAGAGTAGTGCGGCCTCCGAGCTCGC                         AGAAGCACCCACG  -24
SAFB2mut1: AGGGCTAGAGCCGGCAGTTGAGCAGAGTAGTGCGGCCTCCGAGCTCGCGGAGGCCTCTAAGCGAGGAGCTCGCAGAAGCACCCA  +1
SAFB2mut2: AGGGCTAGAGCCGGCAGTTGAGCAGAGTAGTGCGGCCTCCGAGCTCGCGGAGGCCTCTAAGCGAGGAGCTCGCAGAAGCACCCA  +1

ERH(+/-)/SAFB1(+/-)/SAFB2(-/-) mutant#6
ERHwt:  TCTCACACCATTTTGCTGGTACAGCCTACC AAGAGGCCAGAAGGCAGAACTTATGCTGACTACGAATCTGTGAATGAATGCATGGAAG
ERHmut: TCTCACACCATTTTGCTGGTACAGCCTACCCAAGAGGCCAGAAGGCAGAACTTATGCTGACTACGAATCTGTGAATGAATGCATGGAAG  +1
SAFB1mut1: GCTAGAGCCGGCAGTTGAGCAGAGTAGTGCGGCCTCCGAGCTCGCGGAGGCCTCTAAGCGAGGAGCTCGCAGAAGCACCCACG  +1
SAFB1mut2: GCTAGAGCCGGCAGTTGAGCAGAGTAGTGCGGCCTCCGAGCTCGCGGAG                CTCGCAGAAGCACCCACG  -15
SAFB2mut1: AGGGCTAGAGCCGGCAGTTGAGCAGAGTAGTGCGGCCTCCGAGCTCGCGGAGGCCTCTATAGCGAGGAGCTCGCAGAAGCACCCA  +2
SAFB2mut2: AGGGCTAGAGCCGGCAGTTGAGCAGAGTAGTGCGGCCTCCGAGCTCGCGGAGGCCTCTA       GAGCTCGCAGAAGCACCCA  -5

DGCR8(-/-) mutant#9
mut1: GTCTTTACTGCGCATGTATGGCCGTGAGAGCA   AAGATGGTCAAGCAG  -2
mut2: GTCTTTACTGCGCATGTATGGCCGTGAGAGCAGCAAAGATGGTCAAGCAG  +1
```

Yellow highlighted sequence is the NGG motif for Cas9 recognition. Underlined sequences containing pre-mir-3618 were deleted.

```
pre-mir-3618 deletion in WT HEK293T cells mutant#29
mut1: GCCAGTCACTTAAGCTGAGTGCATTGTGATTTCCAATAATTGAGGCAGTGGTTCTAAAAGCTGTCTACATTAATGAAAAGAGCAATGTGGCCAGCTTGACTAA
mut2: GCCAGTCACTTAAGCTGAGTGCATTGTGATTTCCAATAATTGAGGCAGTGGTTCTAAAAGCTGTCTACATTAATGAAAAGAGCAATGTGGCCAGCTTGACTAA

pre-mir-3618 deletion in SAFB1(+/-)/SAFB2(-/-)#14 HEK293T cells mutant#21
mut1: GCCAGTCACTTAAGCTGAGTGCATTGTGATTTCCAATAATTGAGGCAGTGGTTCTAAAAGCTGTCTACATTAATGAAAAGAGCAATGTGGCCAGCTTGACTAA
mut2: GCCAGTCACTTAAGCTGAGTGCATTGTGATTTCCAATAATTGAGGCAGTGGTTCTAAAAGCTGTCTACATTAATGAAAAGAGCAATGTGGCCAGCTTGACTAA
```

Supplementary Figure 2. Genotyping of *ERH*, *SAFB1/2* and *mir-3618* mutations.
Shown are the sequenced alleles of *ERH*, *SAFB1*, *SAFB2* indels and *mir-3618* deletions recovered in HEK293T cells used in this study.

Shang et al,
Supplementary Figure 2

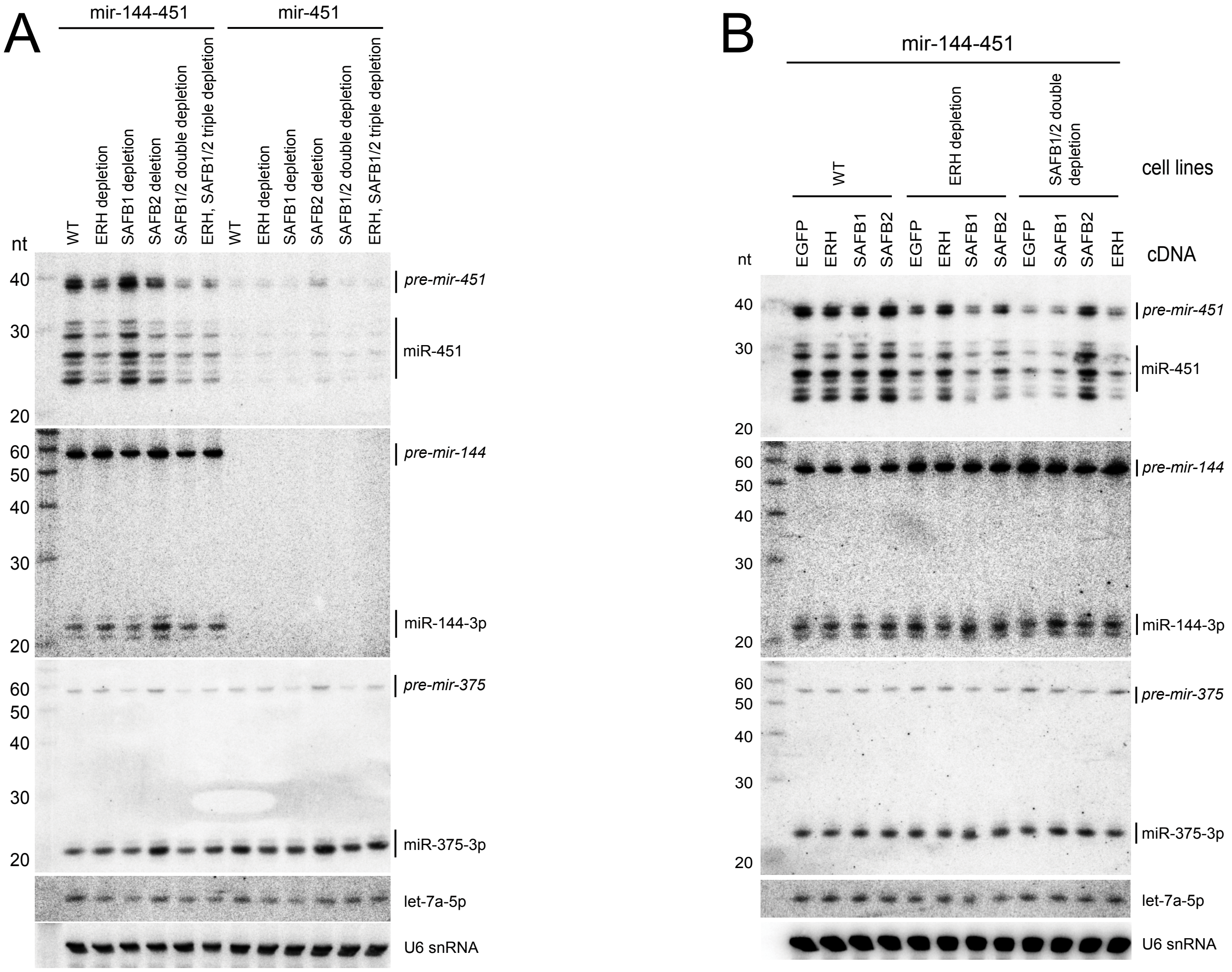


**WT:** wild-type HEK293T cells + shGFP

**ERH depletion:** *ERH[+/-]* cells + shERH

**SAFB1 depletion:** *SAFB1[+/-]* cells + shSAFB1

**SAFB2 deletion:** *SAFB2[-/-]* cells

**SAFB1/2 double depletion:**
*SAFB1[+/-], SAFB2[-/-]* cells + shSAFB1

**ERH, SAFB1/2 triple depletion:**
*ERH[+/-]; SAFB1[+/-], SAFB2[-/-]* cells
+shERH/shSAFB1

Supplementary Figure 3. Processing of *mir-144/451* or solo *mir-451* in cells depleted of Microprocessor cofactors.
(A) Comparison of miRNA processing by Northern blotting from HEK293T cells depleted of either of *ERH*, *SAFB1*, *SAFB2* or their combinations. Co-transfected *mir-375* and endogenous let-7a and U6 snRNAs were probed as controls. RNA size markers (nt) are shown on the left. Microprocessing of suboptimal *pri-mir-451*, but not optimal *pri-mir-144* and *pri-mir-375*, is greatly repressed in ERH or SAFB2 single mutant cells, while no defect was observed in SAFB1 single mutant cells. But interestingly, SAFB1/2 double mutant showed stronger repression on *pri-mir-451* processing than SAFB2 single mutant, indicating a collaborative activity of SAFB1 and SAFB2. In contrast, absence of the neighboring *mir-144* hairpin almost completely repressed mir-451 expression in all these cell lines.
(B) Cross-rescue of *pri-mir-451* processing in mutant cells. While ERH cDNA can restore miR-451 biogenesis, overexpressing SAFB2 also partially restored miR-451 in ERH mutant cells. In contrast, only SAFB2, but not SAFB1 and ERH, can rescue mir-451 biogenesis in SAFB1/2 double mutant cells. These results further confirmed the unique and critical role of SAFB2 in cluster assistance.

Shang et al,
Supplementary Figure 3

A No dimerization of the ERH mutants.

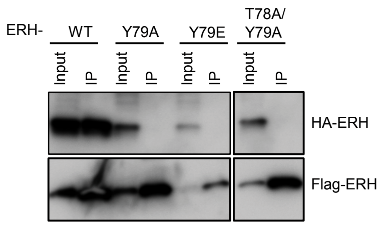


B DGCR8 variants are stable proteins.

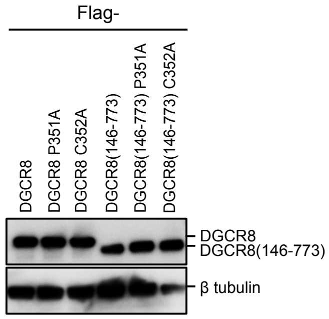


C DGCR8 does not require ERH interaction to promote cluster assistance

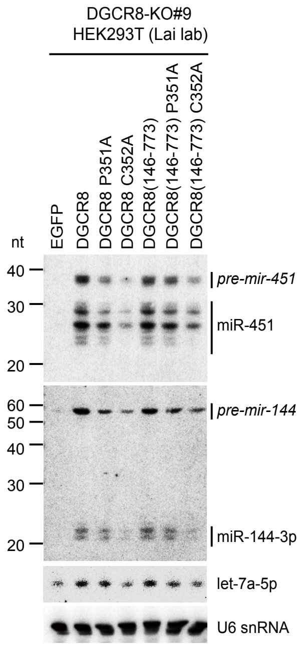


Supplementary Figure 4. Physical and function interactions between ERH and miRNA pathway factors. (A) Co-immunoprecipitation (co-IP) assays of wildtype and variant ERH proteins bearing the designated mutations confirms that these mutants ablate dimerization of ERH. (B) Western blotting of DGCR8 variants lacking the N-terminal ERH interaction domain (residues 1-145) and/or that carry mutations in the DGCR8 dimerization domain confirm that all of these accumulate to similar levels in cells. (C) Northern blotting of rescue assays of DGCR8 variants transfected into *DGCR8-KO* cells. Deletion of the N-terminal domain does not affect rescue capacity of DGCR8 for optimal or suboptimal miRNAs, but loss of dimerization compromises its activity (especially for point mutant C352A). This experiment is similar to main Figure 3H, but uses an independent *DGCR8-KO* line generated in our lab.

Shang et al,
Supplementary Figure 4

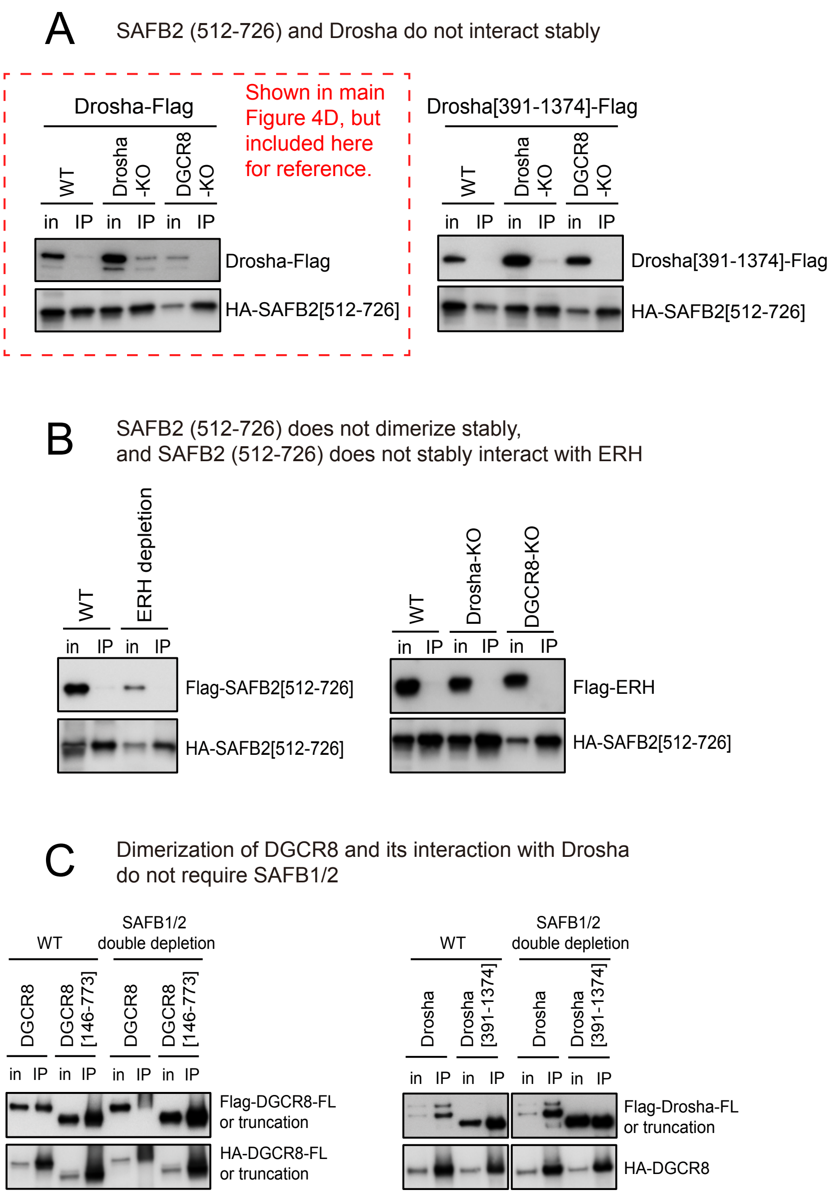


Supplementary Figure 5. Interaction tests between SAFB1/2 and Microprocessor factors.
HEK293T cells were transfected with the indicated tagged plasmids. HA-tagged factors were immunoprecipitated and Flag-tagged factors were tested for co-immunoprecipitation (co-IP). (A) The minimal fragment SAFB2[512-726] fully supports miRNA cluster assistance in SAFB1/2 mutant cells. Co-IP assays showed only mild, or no stable interactions between SAFB2[512-726] and full length Drosha, or Drosha deleted for its N-terminus (reported to contain the SAFB2 binding region). Assays were conducted in wildtype and knockout cells for both Microprocessor components. (B) Co-IP assays did not detect stable dimerization of functional SAFB2[512-726], and did not detect stable association of SAFB2[512-726] with ERH, either in wildtype cells or Microprocessor mutant cells. (C) Co-IP assays indicate that neither dimerization of DGCR8, nor stable formation of Microprocessor complex, requires SAFB1/2.

Shang et al,
Supplementary Figure 5

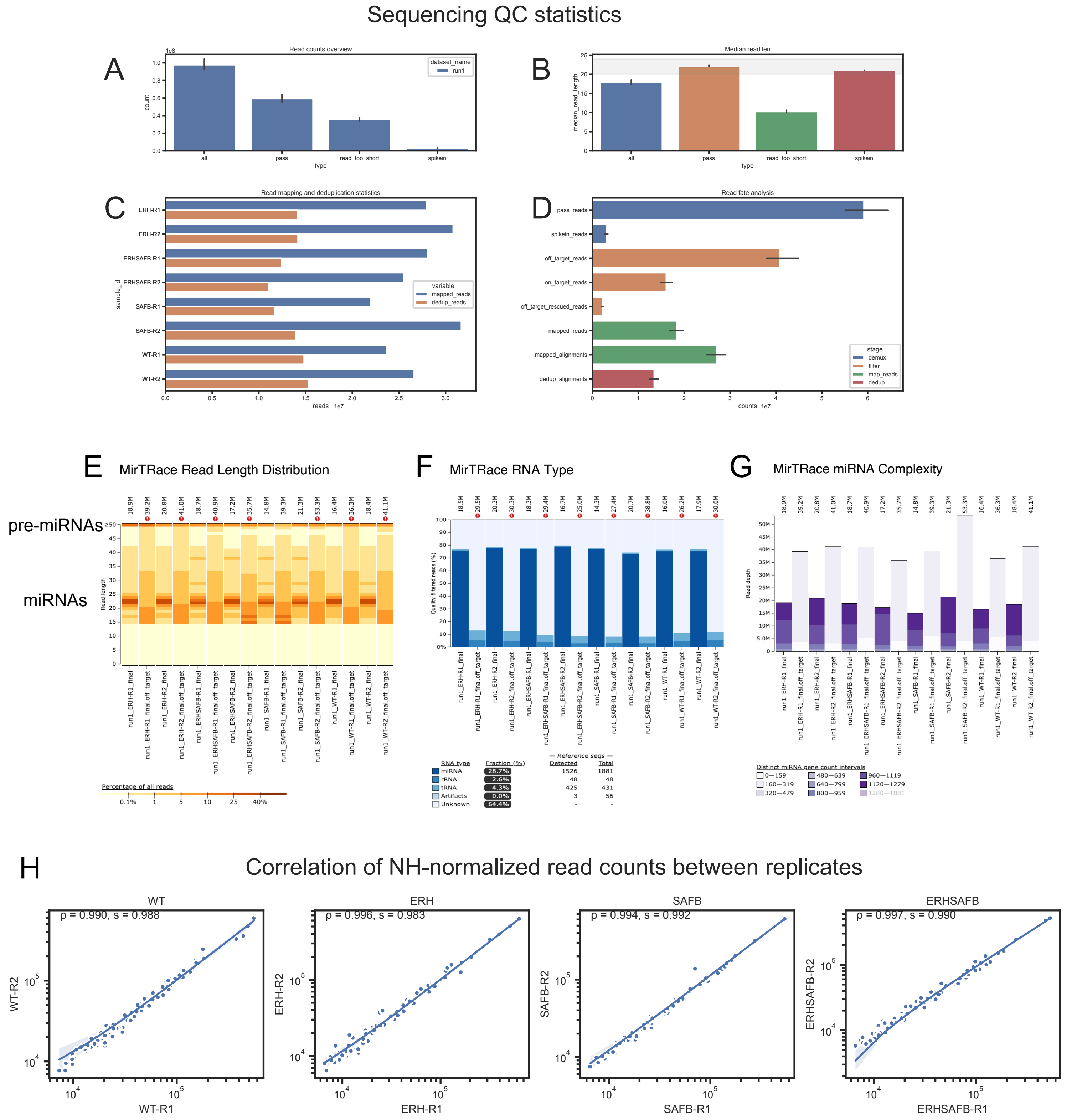


Supplementary Figure 6. Processing and QC statistics of SLAM-seq data from Microprocessor cofactor mutants.

(A) Raw read counts before (all) and after demultiplexing (pass: reads for further processing, spikein: reads assigned to spike-in sequences, read_too_short: filtered reads). (B) Median read lengths of the respective groups (error bar indices 95% ci over all datasets). The grey band highlights the expected size of mature miRNAs (20-24 nts). (C) Deduplication statistics. mapped_reads: number of alignments before deduplication. dedup_reads: number of reads after deduplication. (D) Read fate statistics from our small RNA analysis pipeline, error bars indicate variation between samples. Final miRNA counting was based on deduplicated alignments. (E) Size distribution of SLAM-seq libraries from 6hr labeling of wt, ERH, SAFB1/2 and ERH+SAFB1/2 HEK293T cells. Filtering was done to remove potentially off-target mappings. Of the pass reads ("final" columns), the majority of reads are in the size range of mature miRNAs, but there are also minor populations of pre-miRNA-sized reads. (F) RNA types of reads from the pass and off-target populations. (G) miRNA complexity of the pass and off-target populations. (H) The replicate libraries are well-correlated.

Shang et al,
Supplementary Figure 6

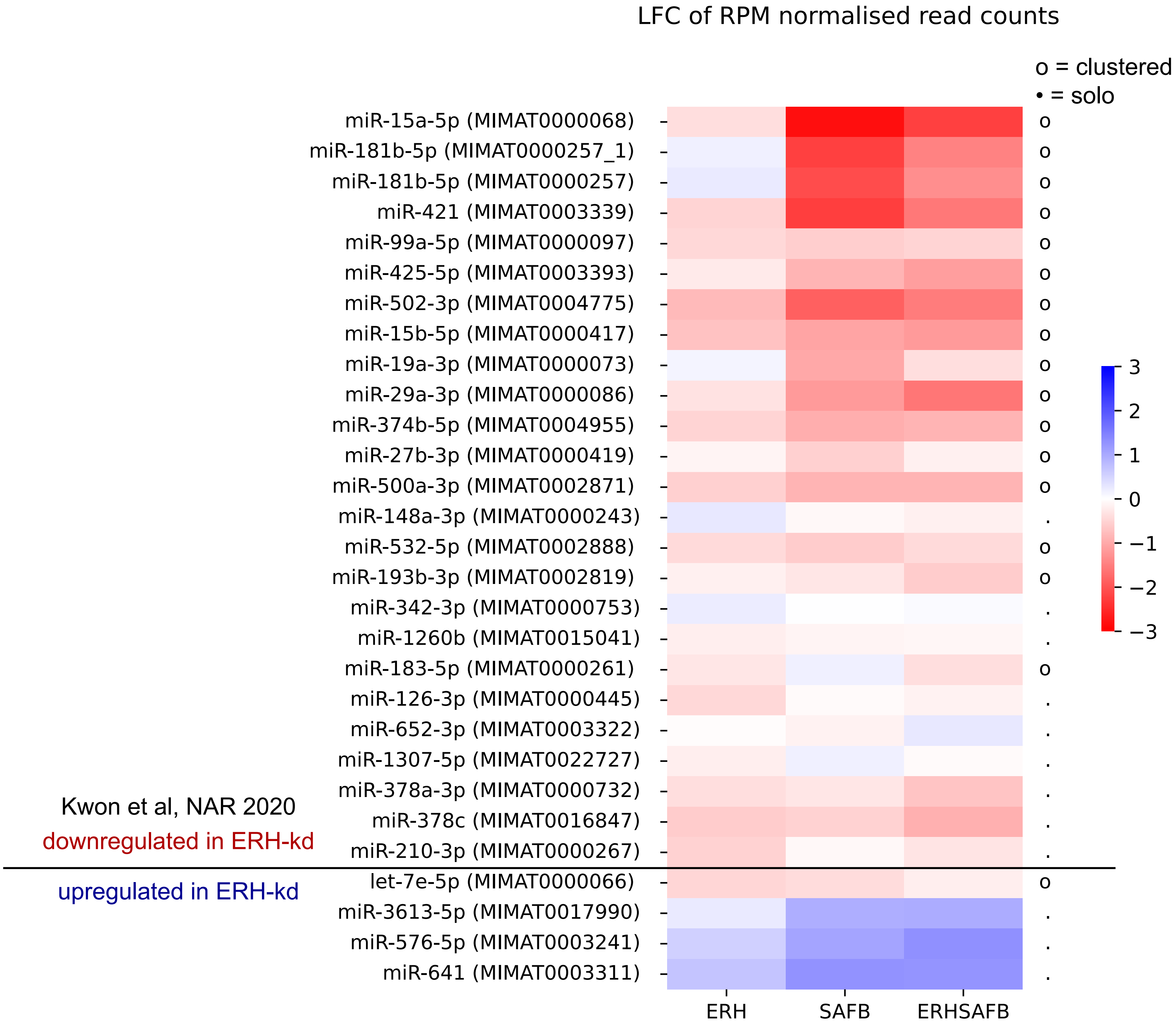


Supplementary Figure 7. Analysis of miRNAs previously assessed for dependency on Microprocessor cofactors.

Shown are the miRNAs reported by Kwon et al Nucleic Acids Research 2020 (Figure 7) to be dependent on ERH and/or SAFB1/2, using different cell models. For ease of comparison, we plotted these in the same order as in the prior study, using our data from ERH, SAFB1/2 and ERH+SAFB1/2 mutant HEK293T cells, and recorded their clustered status with circles or dots (to the right of the heatmap). For the most part, our data agree with the prior study, except that let-7e-5p was mildly downregulated in our data (as opposed to upregulated in ERH-kd), and a few other miRNAs were mildly downregulated in our data (but upregulated in prior ERH-kd data).

Shang et al,
Supplementary Figure 7

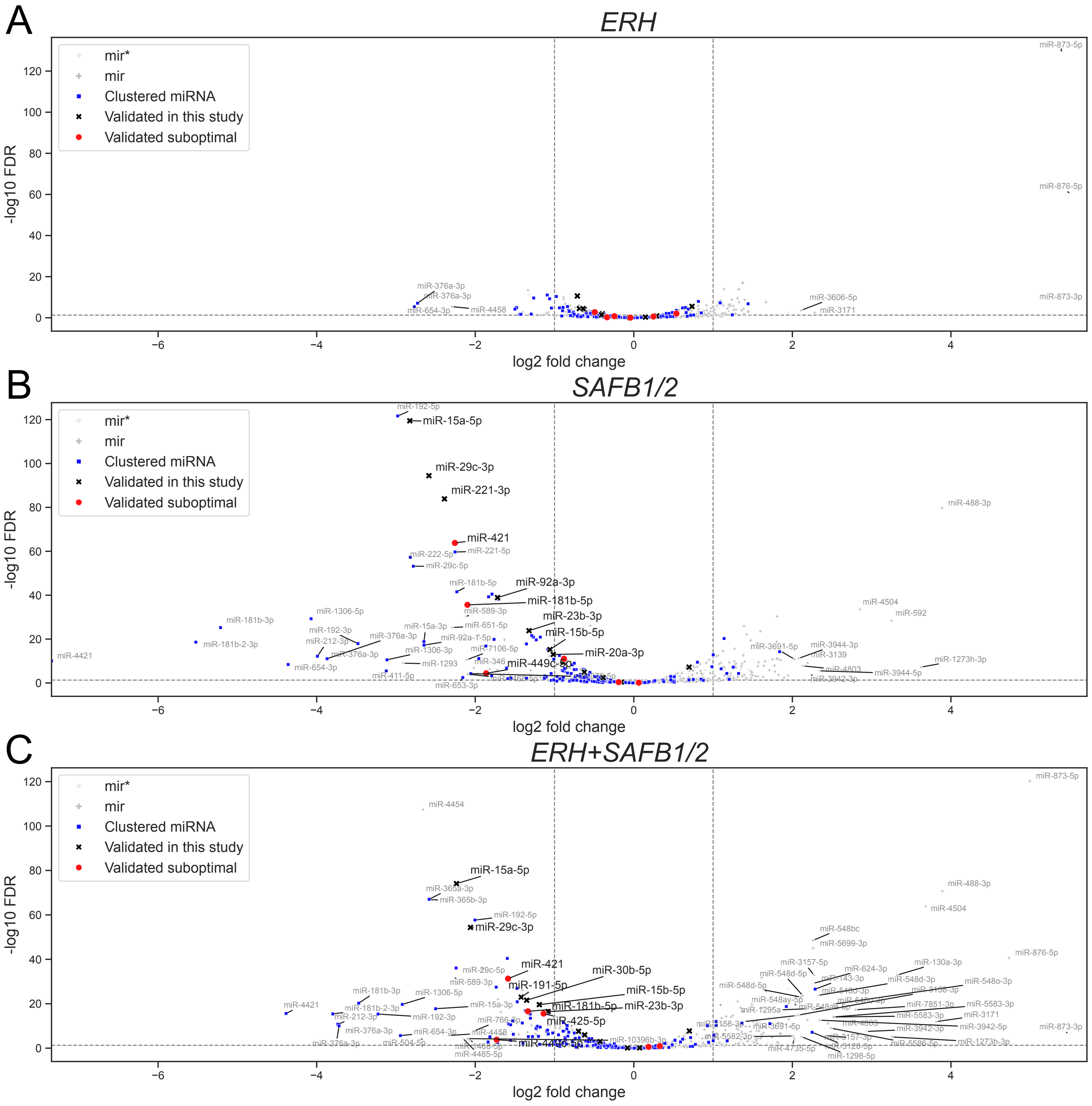


Supplementary Figure 8. Volcano plots of miRNA dysregulation in mutants of Microprocessor cofactors.

Volcano plots from comparing (A) ERH, (B) SAFB1/2 and (C) ERH+SAFB 1/2 to WT (n=713), a negative log2 fold change indicates downregulation in the respective KD). The plot contains all final miRNAs, clustered and validated suboptimal miRNAs are shown using blue and red color respectively, miRNAs that were tested by Northern blotting in this study are highlighted using black color. Significantly DE miRNAs with |LFC|>2 are labeled.

Shang et al,
Supplementary Figure 8

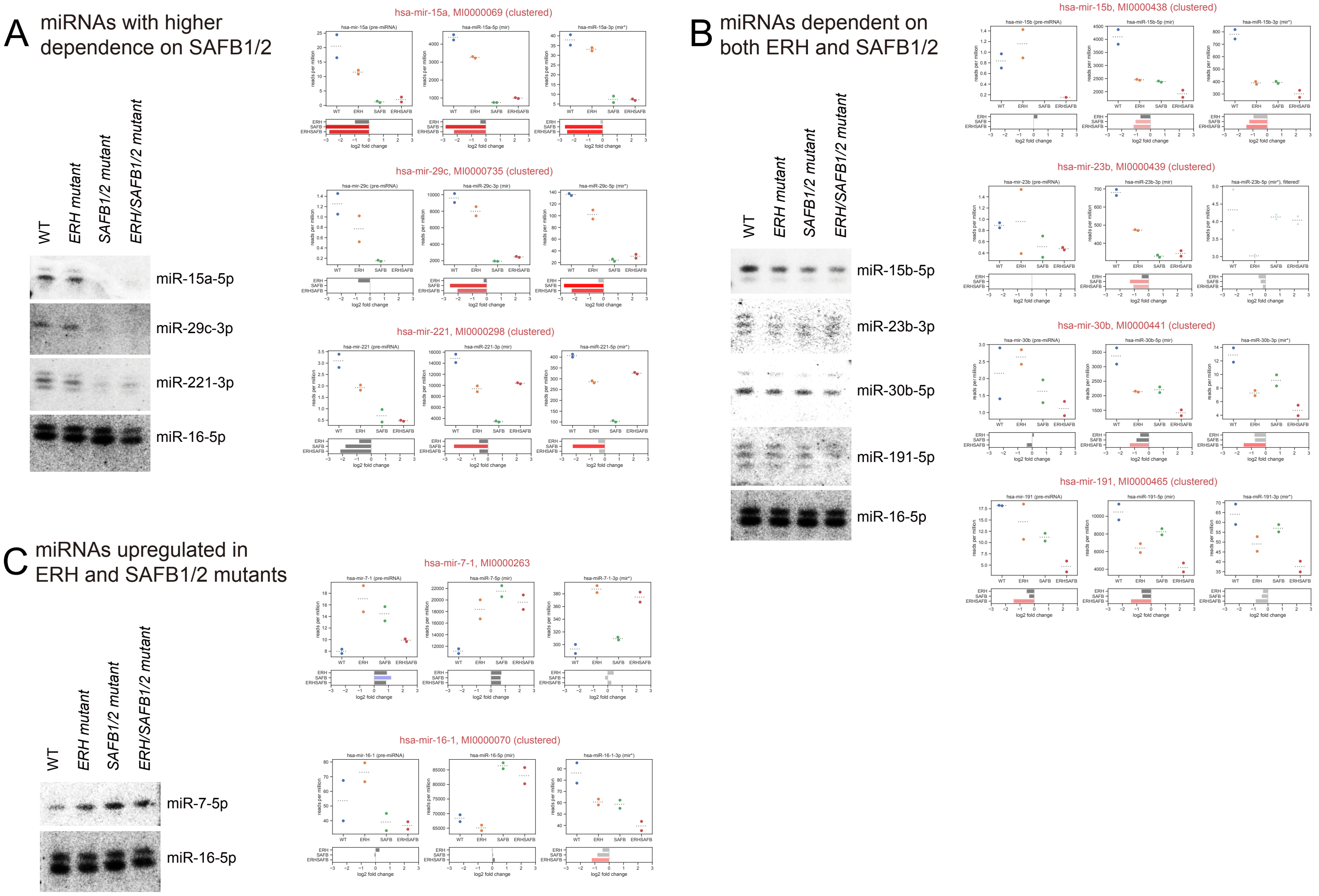


Supplementary Figure 9. Validation of endogenous miRNAs in different genotypes by Northern blots.

Total RNAs of WT, ERH and/or SAFB1/2 mutant HEK293T cells (the same batch of RNAs used for SLAM-seq) were run on three blots to probe different miRNAs that (A) are preferentially downregulated in SAFB1/2 compared to ERH, (B) are dependent on both ERH and SAFB1/2, or that (C) are upregulated in ERH and SAFB1/2 mutants. miR-16 is an optimal miRNA that is relatively unchanged in mutants and deep sequencing, and was used as control.

Shang et al,
Supplementary Figure 9

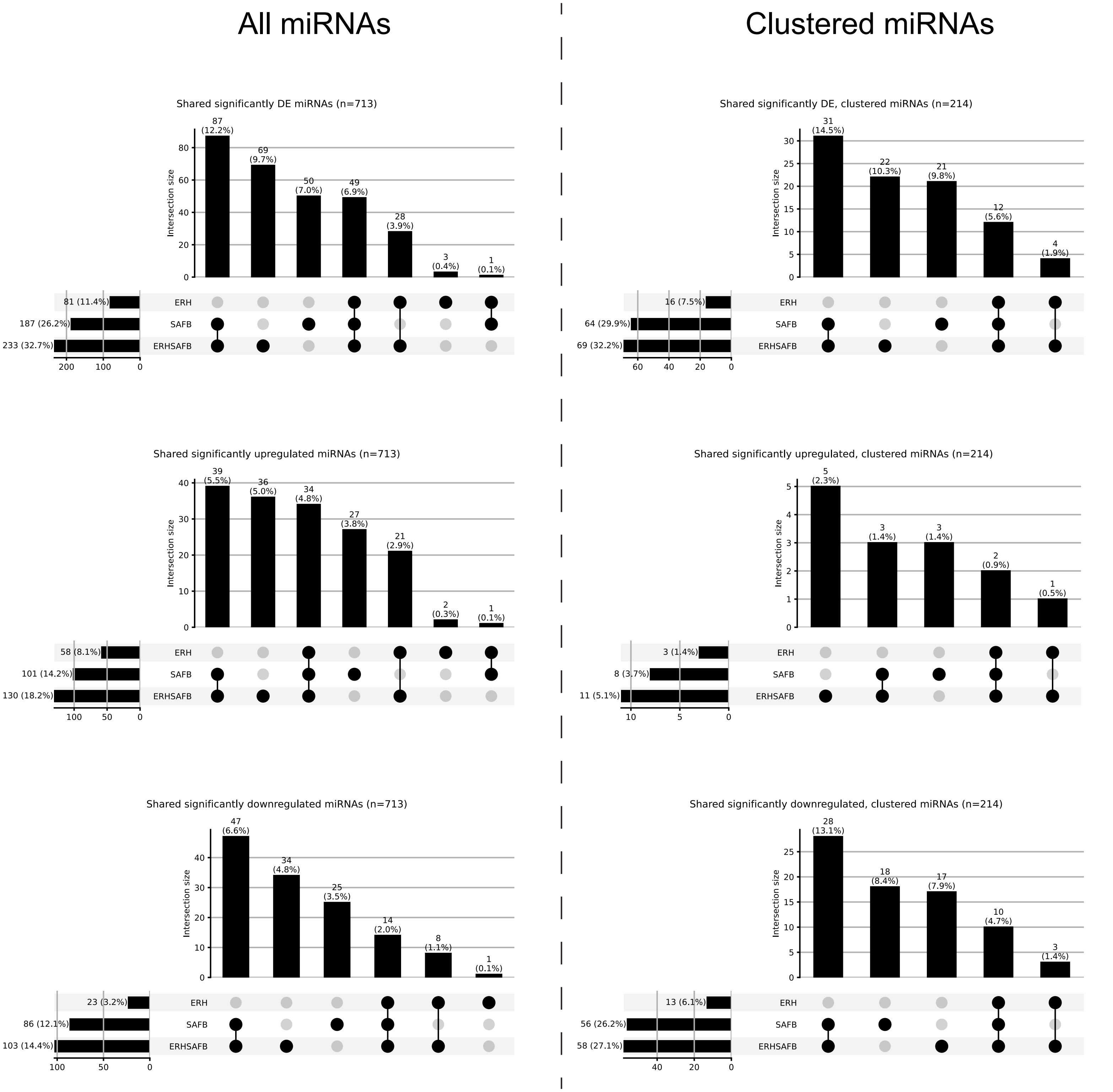


Supplementary Figure 10. UpSet plots of miRNA dysregulation in mutants of Microprocessor cofactors.

Shown are UpSet plots of differentially expressed miRNAs including all (left panels) and only clustered miRNA loci (right panels). These were divided according to plots of all miRNAs (top panels), only upregulated miRNAs (middle panels) and only downregulated miRNAs (bottom panels). It can be seen that SAFB1/2 and ERH+SAFB1/2 mutants account for the majority of downregulated miRNAs, and that a substantial portion of these are also shared by ERH mutants. In addition, amongst clustered miRNAs, there are far more that are downregulated than upregulated in these mutants, consistent with the notion that disruption of cluster assistance is a major drive of miRNA dysregulation in these mutants, as opposed to other direct roles or indirect consequences of these mutant cells.

Shang et al,
Supplementary Figure 10

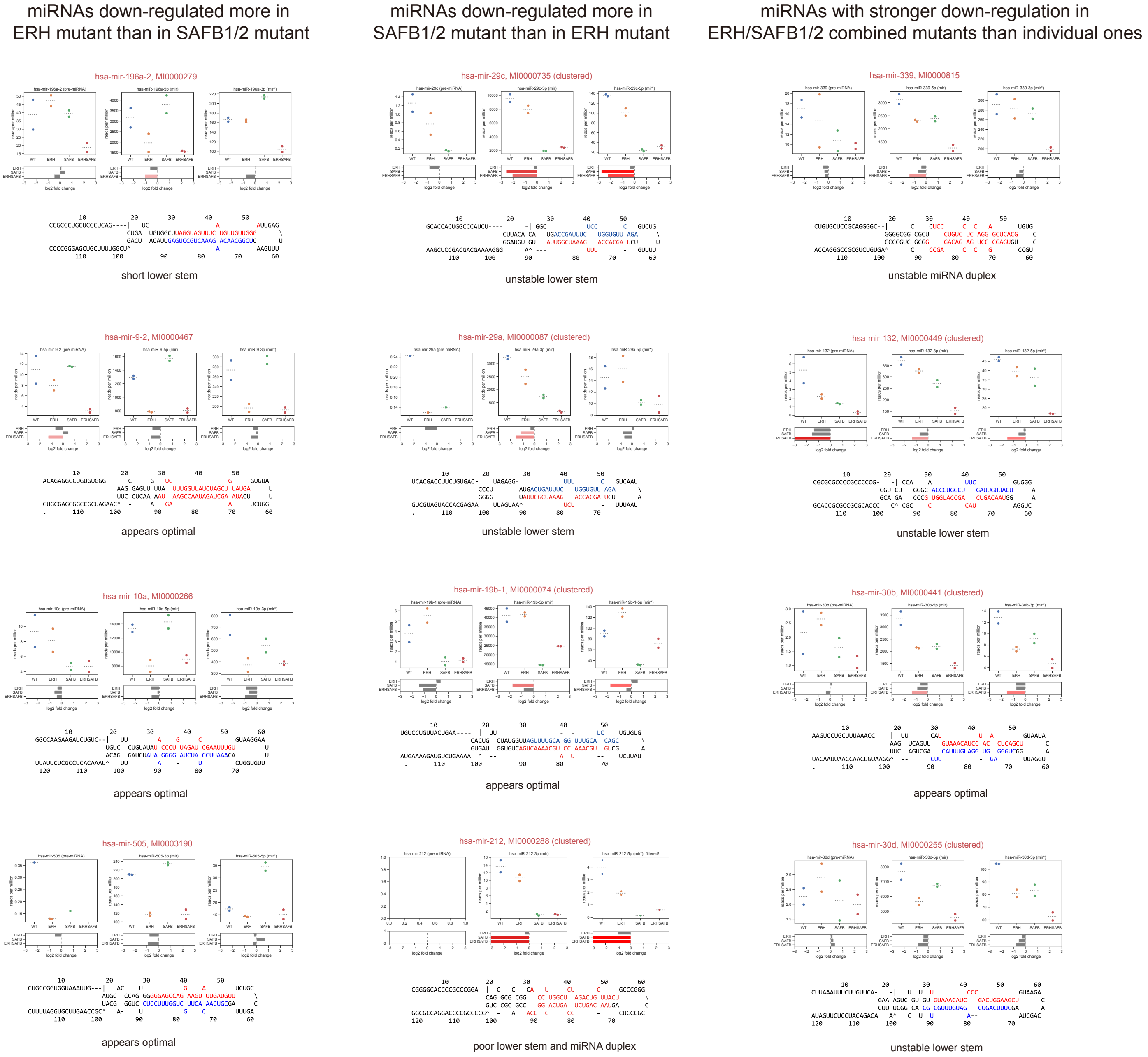


Supplementary Figure 11. Additional examples of suboptimal miRNAs with dependency on ERH and/or SAFB1/2.

We extracted illustrative miRNAs with preferential downregulation in ERH mutant, in SAFB1/2 mutant (without or with additional ERH depletion), or in combined ERH+SAFB1/2 mutant HEK293T cells. Note that as SAFB1/2 depletion had the largest effect on miRNA dysregulation in these datasets, we were able to select examples that show more substantial defects in miRNA expression than in other cases. For example, there are few miRNAs that show an enhanced downregulation in the combined triple mutant data. For all genotypes, we highlighted example loci where similar trends were observed in both mature miRNA and passenger strand (miRNA*) species of the given miRNA hairpin, which helped support the notion that these reflect biogenesis defects (as opposed to stability effects). We show predicted secondary structures for these loci. In many cases, they exhibit predicted subtoptimal stems that may underlie their sensitivity to Microprocessor cofactors. Amongst ERH-sensitive miRNAs, many exhibit seemingly typical pri-miRNA structures. It remains to be see if these indeed hairpin suboptimality that is not reflected by structure predictions, or if these ERH-dependent miRNAs involve effects other than cluster assistance.

Shang et al,
Supplementary Figure 11

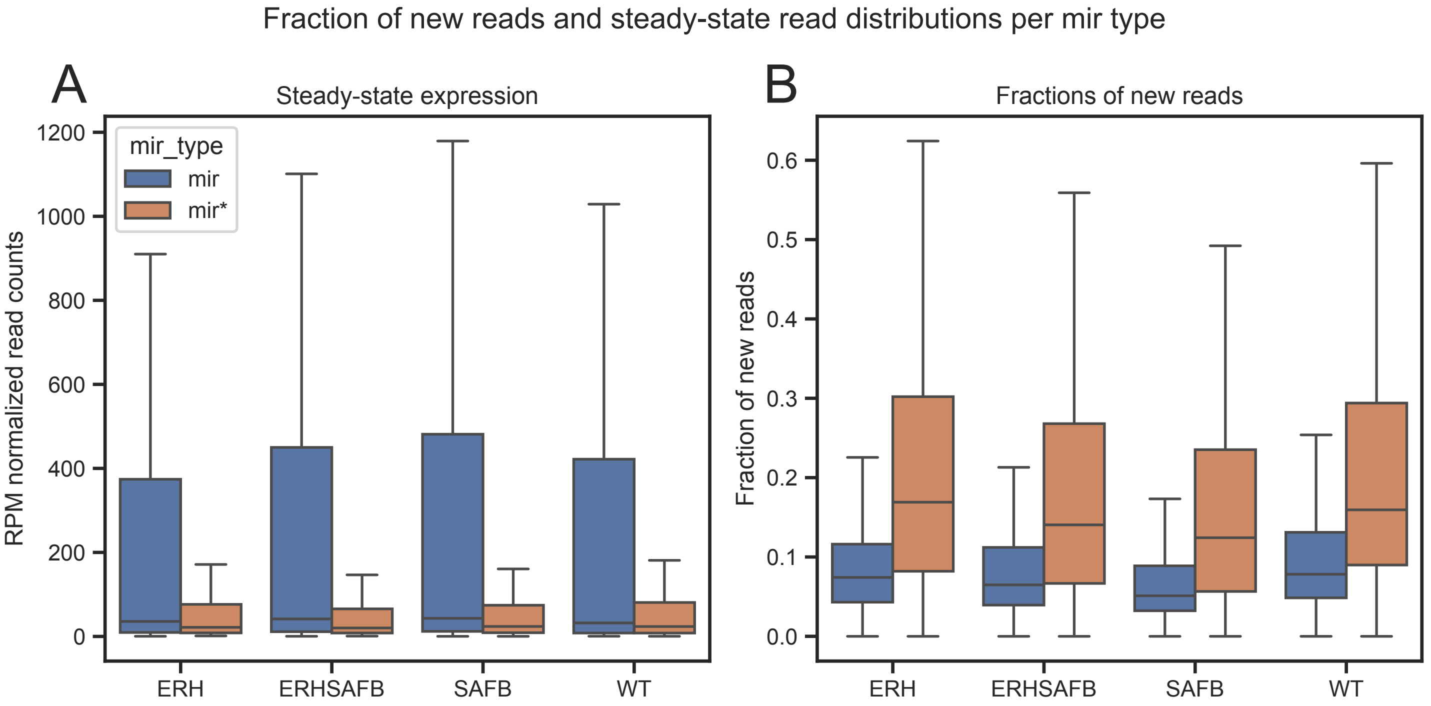


Supplementary Figure 12. Quality control assessment of newly-synthesized miRNAs in SLAM-seq data.

SLAM-seq libraries were constructed following labeling with 4sU for 6 hours. Fractions of reads from newly synthesized miRNAs were estimated from the observed T-to-C conversion patterns as described in Methods. (A) Amongst total reads, mature miRNA strands dominate all the libraries for the four genotypes. (B) However, amongst newly-synthesized reads, miRNA* (passenger strands) are heavily enriched. This behavior is expected, since miRNA* species are degraded following their ejection from the mature Argonaute-miRNA complex. This assessment supports that the SLAM-seq protocol indeed captured newly-synthesized small RNAs (transcribed within the 6 hr labeling period).

Shang et al,
Supplementary Figure 12

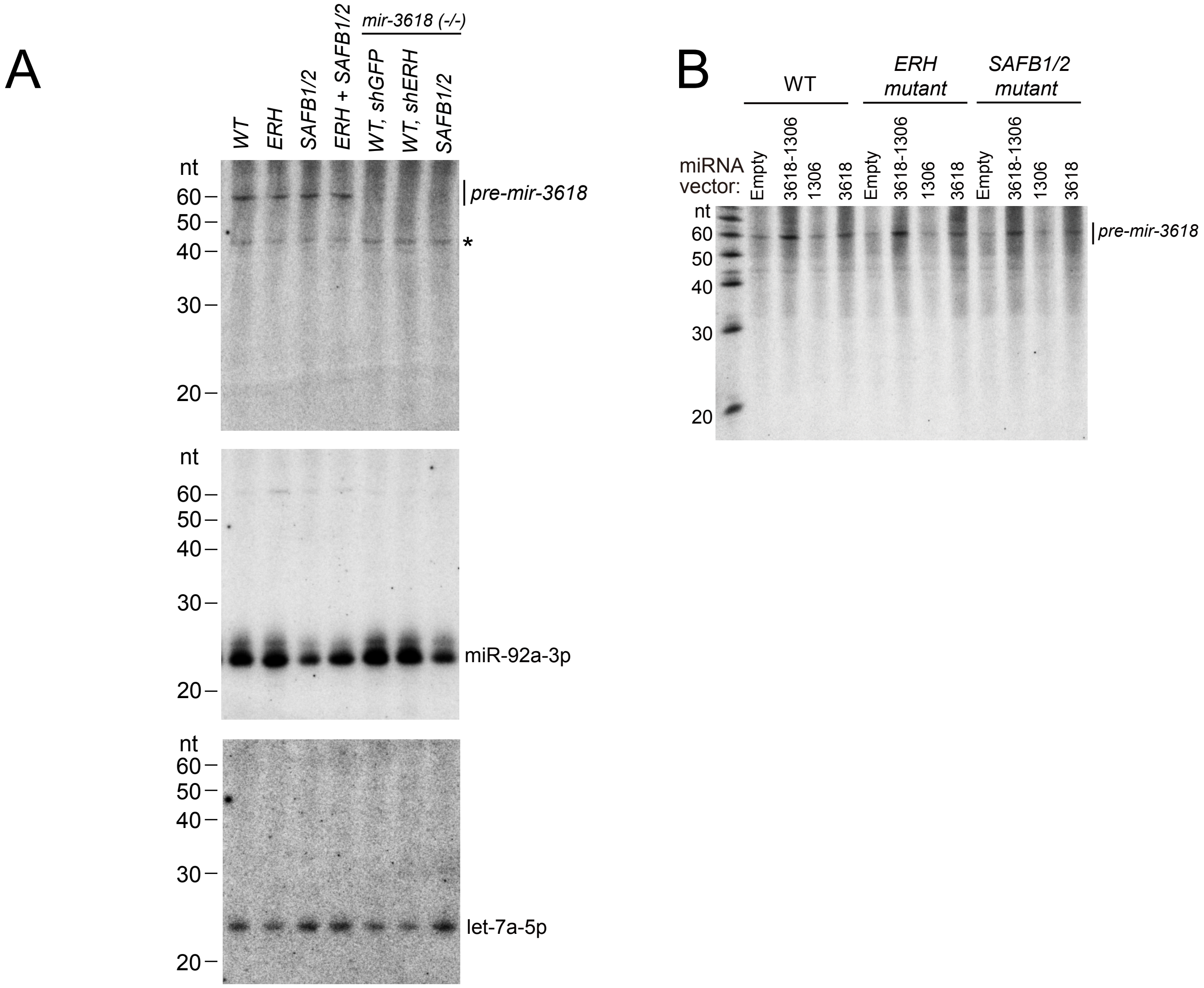


Supplementary Figure 13. Additional Northern blotting of *DGCR8* miRNAs.
(A) Northern blotting shows that endogenous *pre-mir-3618* is readily detected, and is lost in *mir-3618* deletion HEK293T cells. However, no mature miR-3618 is visible. Other probings show mature miR-92a-3p and let-7a-5p. Note that these are the same original blots that are cropped in main Figure 7C to highlight the specific bands. (B) Ectopic expression of DGCR8 miRNA constructs containing *mir-3618*, *mir-1306*, or both, increase signals for *pre-mir-3618*, but still do not reveal mature miR-3618.

Shang et al,
Supplementary Figure 13
